# Spatiotemporal landscape of SARS-CoV-2 pulmonary infection reveals *Slamf9*^+^*Spp1*^+^ macrophages promoting viral clearance and inflammation resolution

**DOI:** 10.1101/2022.05.03.490381

**Authors:** Boyi Cong, Xuan Dong, Zongheng Yang, Pin Yu, Yangyang Chai, Jiaqi Liu, Meihan Zhang, Yupeng Zang, Jingmin Kang, Yu Feng, Yi Liu, Weimin Feng, Wei Deng, Fengdi Li, Qinyi Yu, Yan Gu, Zhiqing Li, Shuxun Liu, Xun Xu, Nanshan Zhong, Xianwen Ren, Chuan Qin, Longqi Liu, Jian Wang, Xuetao Cao

## Abstract

While SARS-CoV-2 pathogenesis has been intensively investigated, the host mechanisms of viral clearance and inflammation resolution are still elusive because of the ethical limitation of human studies based on COVID-19 convalescents. Here we infected Syrian hamsters by authentic SARS-CoV-2 and built an ideal model to simulate the natural recovery process of SARS-CoV-2 infection from severe pneumonia^1,2^. We developed and applied a spatial transcriptomic sequencing technique with subcellular resolution and tissue-scale extensibility, *i.e.*, Stereo-seq^3^, together with single-cell RNA sequencing (scRNA-seq), to the entire lung lobes of 45 hamsters and obtained an elaborate map of the pulmonary spatiotemporal changes from acute infection, severe pneumonia to the late viral clearance and inflammation resolution. While SARS-CoV-2 infection caused massive damages to the hamster lungs, including naïve T cell infection and deaths related to lymphopenia, we identified a group of monocyte-derived proliferating *Slamf9*^+^*Spp1*^+^ macrophages, which were SARS-CoV-2 infection-inducible and cell death-resistant, recruiting neutrophils to clear viruses together. After viral clearance, the *Slamf9*^+^*Spp1*^+^ macrophages differentiated into *Trem2*^+^ and *Fbp1*^+^ macrophages, both responsible for inflammation resolution and replenishment of alveolar macrophages. The existence of this specific macrophage subpopulation and its descendants were validated by RNAscope in hamsters, immunofluorescence in hACE2 mice, and public human autopsy scRNA-seq data of COVID-19 patients. The spatiotemporal landscape of SARS-CoV-2 infection in hamster lungs and the identification of *Slamf9*^+^*Spp1*^+^ macrophages that is pivotal to viral clearance and inflammation resolution are important to better understand the critical molecular and cellular players of COVID-19 host defense and also develop potential interventions of COVID-19 immunopathology.

## Introduction

COVID-19, caused by severe acute respiratory syndrome coronavirus 2 (SARS-CoV-2) infection, has become a pandemic worldwide^4,5^. While most patients develop mild, moderate to self-limiting symptoms, severe COVID-19 could result in death from severe pneumonia, acute respiratory distress syndrome (ARDS) and multi-organ failure^6,7^. Multiple studies have analyzed the innate and adaptive immune profiles of multiple tissues and blood samples of COVID-19 patients, shedding light on the immunological and immunopathological mechanisms for inflammation initiation and tissue damage^8–14^. However, it is still elusive how SARS-CoV-2 infection was dynamically cleared and how severe pneumonia was finally resolved because of the difficulty of human studies.

Identifying cell subpopulations responsible for clearing SARS-CoV-2 or resolving pulmonary inflammation is pivotal to understanding COVID-19 pathogenesis and developing antiviral therapeutics^15^. To conquer the research difficulty, here we established an ideal model to simulate the viral clearance and inflammation resolution process by infecting Syrian hamsters with appropriate titers of authentic SARS-CoV-2. Syrian hamsters, naturally susceptible to SARS-CoV-2 infection, have been widely used in hundreds of studies to mimic severe COVID-19 with the aim of understanding the pathogenesis of SARS-CoV-2 infection, developing and evaluating the effectiveness of new drugs, vaccines and antibodies^1,2,16–19^. However, no reports have been published about the spatiotemporal landscape of SARS-CoV-2 clearance together with inflammation resolution and tissue repair. Here we applied our recently developed Stereo-seq^3^ and droplet-based high-throughput scRNA-seq to the lungs of 45 hamsters infected with or without authentic SARS-CoV-2. A comprehensive spatiotemporal cellular and molecular atlas of the lung tissues was generated, covering whole lung lobes and the whole process from tissue damage and severe pneumonia to viral clearance and inflammation resolution. The data illustrated that a group of *Slamf9*^+^*Spp1*^+^ macrophages in the lung, which were specifically induced from monocytes during the acute infection phase and also kept actively proliferating, could recruit neutrophils to collaboratively clear SARS-CoV-2 from hamster lungs. After viral clearance, *Slamf9*^+^*Spp1*^+^ macrophages differentiated into *Trem2*^+^ and *Fbp1*^+^ macrophages to resolve inflammation, replenish alveolar macrophages, and repair damaged lungs. The data resource of the spatial dynamics of SARS-CoV-2 infection and the identification of *Slamf9*^+^*Spp1*^+^ macrophages for SARS-CoV-2 clearance and inflammation resolution provide valuable information for controlling the current COVID-19 pandemic.

## Results

### Whole-lung single-cell spatiotemporal changes of SARS-CoV-2 infected hamsters

To map the panoramic view of SARS-CoV-2 lung infection and clearance, we obtained a comprehensive cellular and molecular atlas for the spatiotemporal changes of hamster lungs before and after authentic SARS-CoV-2 infection (**Fig. 1a**). Transcriptomic data at single-cell and subcellular resolution (with 220 nm spots and 500 nm distance between spots) were achieved for centimeter-scale lung tissues across two weeks (∼1 cm^2^, almost covering the entire lung lobe of hamsters at a certain cross-section). Two types of SARS-CoV-2 titers with 10-fold differences were applied to intranasally infect hamsters so that the pathological changes of SARS-CoV-2-induced severe pneumonia and particularly the viral clearance and inflammation resolution process could be naturally observed and dynamically investigated (Fig. 1b,c**, Supplementary Table 1**).

**Fig. 1.**
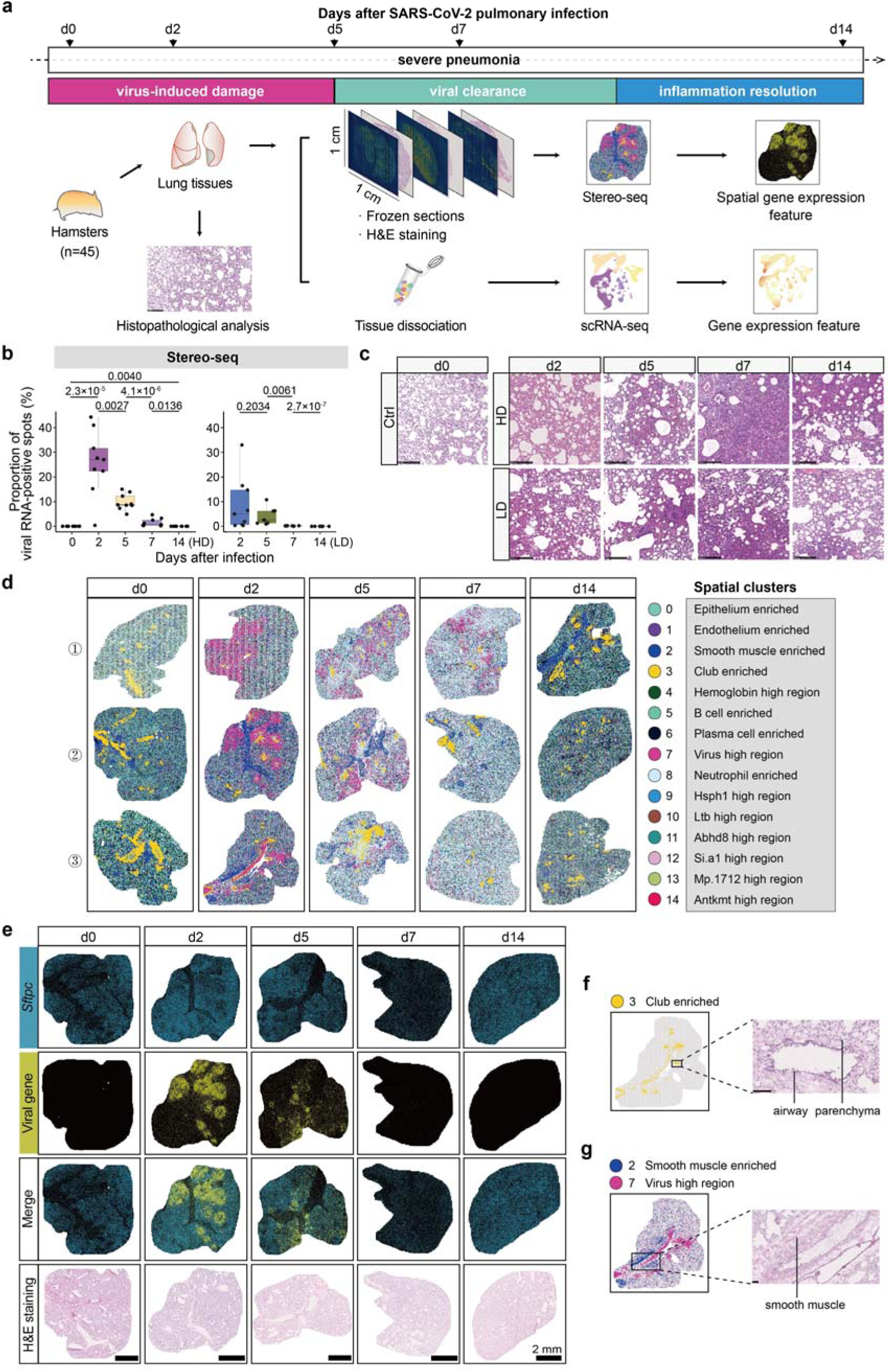
A spatiotemporal landscape of hamster lungs during SARS-CoV-2 infection. **a.** Overview of the experimental design and analysis. **b.** Proportion of SARS-CoV-2 RNA-positive spots (viral count > 0) of nine lung sections at each timepoint (boxplots: center line, median; box bounds, first and third quartiles; whiskers, 1.5 times the interquartile range). Two-sided Student t-test. **c.** The histopathological changes of Ctrl and hamster lungs before and after SARS-CoV-2 infection at different time points (H&E staining, d2, d5, d7 and d14). Scale bar, 250 μm. **d.** Unsupervised clustering of spatial transcriptomic sections identified 15 spatial clusters. **e.** Spatial transcriptomics data of representative hamster lung tissue sections. ATII marker gene *Sftpc*, SARS-CoV-2 genes, and H&E staining of adjacent frozen section were shown. Scale bar, 2000 μm. **f-g.** The highlight of spatial cluster 3 representing club cell-enriched region (**f**), and spatial cluster 2 and 7 representing smooth muscle cell-enriched region and viral genes-highly expressed region (**g**) on a representative section, and the magnified H&E staining image of the framed region. Scale bar, 100 μm.

We obtained spatial transcriptomic data for 81 centimeter-scale lung tissues and scRNA-seq data for 269,199 single cells from 45 hamsters composed of three groups (Ctrl: healthy control; HD: high dose of SARS-CoV-2 titers, 10^5^ TCID50; and LD: low dose of SARS-CoV-2 titers, 10^4^ TCID50), covering five timepoints across two weeks (Fig. 1a**, Extended Data Fig.1a**). These data captured the panoramic spatiotemporal changes of hamster lungs before SARS-CoV-2 infection (d0), with acute tissue damages (d2), demonstrating acute immune responses (d5), exhibiting viral clearance (d7), and approaching inflammation resolution and tissue repair (d14).

To gain an unbiased global overview of cellular and molecular dynamics during SARS-CoV-2 infection at the whole-lung level, we performed unsupervised clustering and identified 15 spatial clusters with distinct gene expression and histological patterns (Fig. 1d**, Supplementary Table 2**). Pulmonary Type II alveolar epithelial cells (ATII, *Sftpc*^+^) dispersed across lung tissues and were enriched in all spatial clusters (Fig. 1e**, Extended Data Fig.1b**). In contrast, club cells (*Scgb3a2*^+^) were mainly detected in spatial cluster 3 and localized surrounding the airway as expected (Fig. 1f**, Extended Data Fig.1b**). The *N* gene of SARS-CoV-2 was highly expressed in spatial cluster 7 with neutrophils infiltrated, which accounted for the main proportion at d2 (**Extended Data Fig.1b,c**). SARS-CoV-2 genes appeared to be dispersed across the entire lung lobe, and the dense regions to be speckled and aggregated alongside the airway indicated by bronchial smooth muscle cells (*Myl9*^+^) in spatial cluster 2 (Fig.1g, **Extended Data Fig.1b**). Such spatial characteristics of SARS-CoV-2 in hamster lungs were further supported by quantitative analysis and additional slides (**Extended Data Fig.1d**). Spatial cluster 5 and 6 were enriched by B cells (*Ighm*^+^) and plasma cells (*Jchain*^+^), respectively, which decreased after infection and partially recovered at d14 (**Extended Data Fig.1b,c,e**), consistent with the clinical observation of lymphopenia and impaired adaptive response at the early stage of COVID-19. In contrast, neutrophils were mainly enriched in d5 and d7 (**Extended Data Fig.1b,c,f**), which might account for severe pneumonia during the acute inflammation stage. In general, these data provide an elaborate map of the spatiotemporal changes of hamster lungs before and after SARS-CoV-2 infection.

Based on the spatial expression of specific marker genes, we calculated the distance between multiple cell types and viral spots according to classical markers defined by literature (**Supplementary Table 3**), and revealed that the distance of viral spots to ATII and club cells was less than 7.5 μm at d2 and d5 (**Extended Data Fig.1g**). Since 7.5 μm is smaller than the diameters of most single cells, such close distance between viral spots and ATII and club cells indicated that ATII and club cells may be the major targets of SARS-CoV-2 acute infection. Likewise, neutrophils were also observed to have a close distance to those viral spots, especially at d5 and d7 (**Extended Data Fig.1g**), suggesting neutrophil phagocytosis of SARS-CoV-2-infected cells or direct recognition of SARS-CoV-2 in the elimination of invading viruses.

The single-cell transcriptomic data revealed 76 cell subpopulations according to unsupervised clustering and manual curation, covering non-immune cells (epithelium, endothelium, fibroblasts and erythrocytes), T/NK/NKT cells, B cells, myeloid cells (alveolar macrophages, AMs; interstitial macrophages, IMs; monocytes/macrophages; dendritic cells, DCs; megakaryocytes), neutrophils, and ILC2 (**Extended Data Fig.2a, Supplementary Table 4**). These cell populations together with SARS-CoV-2 demonstrated different dynamics of lung accumulation during the infection process (**Extended Data Fig.2b-e, Supplementary Table 5**). Multiple cell types demonstrated a decreasing trend from d2 to d7, and then partially restored at d14, which were consistent with the observations of tissue damage and then alleviation. For examples, T and B cells were depleted after acute infection and then partially restored from d7 to d14 (**Extended Data Fig.2d, 3a**), indicating SARS-CoV-2 infection could impair the subsequent induction of adaptive response. The lung-resident AMs are critical innate pulmonary sentinels for respiratory pathogens, playing important roles in homeostasis maintenance and pathogen clearance. Similarly, lung-resident AMs started to decrease after d2, and partially restore at d14 (**Extended Data Fig.2d, 3a**), indicating that SARS-CoV-2 infection damaged the lung-resident AMs to escape lung innate immune defense. In contrast, monocytes/macrophages and neutrophils were enriched at d5 and d7, exhibited huge phenotypic heterogeneity, and demonstrated profound temporal changes (**Extended Data Fig.2d, 3a**), consistent with the spatial heterogeneity revealed by our Stereo-seq data (**Extended Data Fig.1f**). Such spatiotemporal characteristics of B, T, monocytes/macrophages and neutrophils highlighted their potential roles in SARS-CoV-2 clearance, inflammation resolution and tissue repair dynamics. The accompanying great phenotypic heterogeneity of these cells also highlighted the complexity of the whole process from SARS-CoV-2-induced severe pneumonia to inflammation resolution, for which Stereo-seq together with scRNA-seq mapped the cellular and molecular events with subcellular resolution at the tissue scale.

In addition to epithelial cells previously regarded as the main host cells of Sarbecoviruses^20^, our study revealed that diverse pulmonary cell populations were also SARS-CoV-2 RNA-positive (**Extended Data Fig.3b,c**). Active subgenomic transcription of SARS-CoV-2 was indicated in multiple cell types, in which increasing detection rates along the genome of SARS-CoV-2 were observed from 5’ to 3’ (**Extended Data Fig.3d**). Dramatic temporal changes were observed for SARS-CoV-2 RNA-positive cells, which peaked at d2, decreased from d5 to d7, and almost diminished at d14 (**Extended Data Fig.2b,c**). Despite of such global trend, the detection rates of SARS-CoV-2 RNA varied among cell populations. As expected, a high proportion and expression of viral RNA was observed in epithelial cells (**Extended Data Fig.3c**). However, AMs, IMs, monocytes/macrophages, neutrophils, and non-immune cells including endothelial cells and fibroblasts were also characterized by the high subgenomic transcription of SARS-CoV-2 (**Extended Data Fig.3d**), with the detection rates similar to or even exceeding those in epithelial cells. DCs, T and B cells harbored a relatively lower detection rate, particularly in plasma cells (**Extended Data Fig.3d**). These data suggested that SARS-CoV-2 results in wide intra-host transmission across different cell types including AMs, neutrophils, naïve T cells and other pulmonary immune cells, potentially infects these cells to induce cell deaths or other phenotypes, leads to lymphopenia, super-inflammation, profound tissue damages and severe pneumonia, and also triggers host responses to resolve inflammation and tissue repair at the late stage.

### SARS-CoV-2 infection of lung-resident macrophage subpopulation and the accompanying cell deaths

While alveolar epithelial cells are assumed to be the host cells of SARS-CoV-2^21^, which was also confirmed in this study, our data suggested that AMs and other lung-resident immune cells also constituted important populations of SARS-CoV-2 RNA-positive cells (**Extended Data Fig.2e**). Among the 16 subpopulations of macrophages identified in this study, which demonstrated distinct temporal enrichment characteristics, a population of *Marco^+^Mrc1^+^Pparg^+^Ccr2*^-^ lung-resident AM (AM-Cd36, cluster 16) was enriched in d0 and d2, which possessed the highest proportion of virus-positive cells at d2 according to scRNA-seq data (up to 50%, much higher than 26% in epithelial cells; Fig. 2a-c**, Extended Data Fig.3c**). This subpopulation was also characterized by active subgenomic transcription of SARS-CoV-2 implied by the increasing detection rates of SARS-CoV-2 genes along the genome from 5’ to 3’ (Fig. 2d), indicating active SARS-CoV-2 transcription. The infection status of AM-Cd36 was further supported by the transcriptional alterations observed between virus-positive and virus-negative cells in AM-Cd36, of which genes related to “response to virus” (such as *Mx1* and *Isg15)* and “apoptosis” (such as *Bax* and *Foxo3*) were upregulated (Fig. 2e,f**, Supplementary Table 6**). The death fate of AM-Cd36 after SARS-CoV-2 infection indicated by upregulated *Bax* and *Foxo3* was confirmed by the depletion of this type of cells at d5 and d7 and co-localization of the marker genes of AM-Cd36 and genes encoding cell deaths according to Stereo-seq data (Fig. 2g,h).

**Fig. 2.**
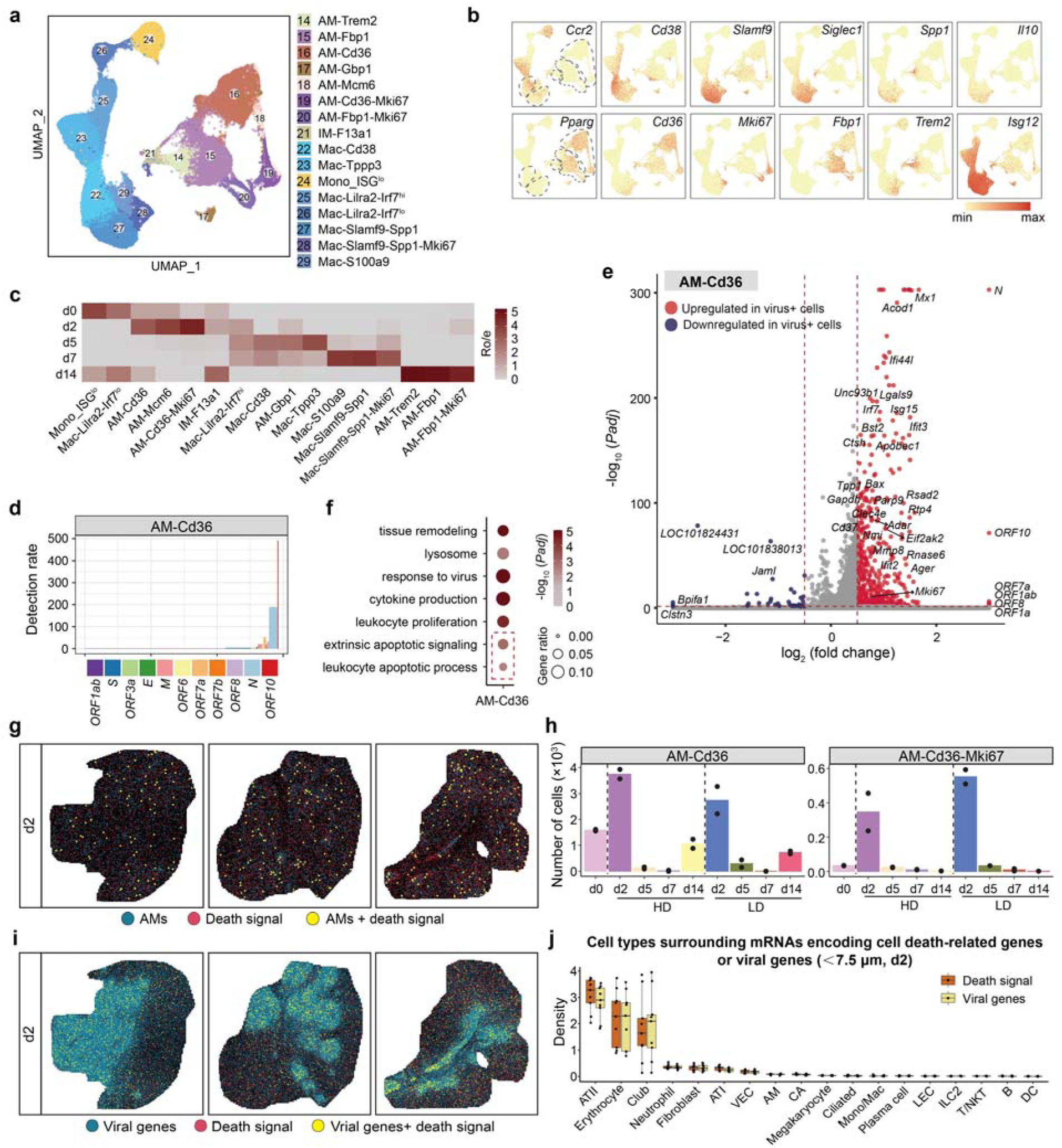
SARS-CoV-2 infection causes massive deaths of lung-resident cells at the early stage. **a.** UMAP projection of AM and mono/mac showing 15 cell subpopulations. **b.** The expression of selected genes defining key AM and mono/mac subpopulations indicated by dashed areas. Gene description: *LOC101838467*, *Isg12*. **c.** Temporal distribution of macrophage subpopulations. Ro/e >1, enrichment; Ro/e<1, depletion. **d.** Detection rates of SARS-CoV-2 genes in AM-Cd36. **e.** Volcano plot of differentially expressed genes between virus-positive and virus-negative cells in AM-Cd36 (cutoff: (|Log_2_FC|>0.5, *Padj*<0.05). **f.** Gene ontology (GO) terms enriched in upregulated genes in virus-positive AM-Cd36. **g.** Spatial distribution of cell death-related genes (death signal) and AMs, represented by three lung sections at d2 (HD group). Magnified size of sparse spots was applied for clarity. **h.** Number of total cells in AM-Cd36 and AM-Cd36-Mki67 detected at each timepoint. **i.** Spatial distribution of virus and death signals, represented by three lung sections at d2 (HD group). Magnified size of sparse spots was applied for clarity. **j.** Density of different cell types in the neighborhood of death signals of nine lung sections at d2 (HD group). Neighborhood: <7.5 μm. Boxplots: center line, median; box bounds, first and third quartiles; whiskers, 1.5 times the interquartile range.

It is worth noting that Stereo-seq data and scRNA-seq data revealed different patterns regarding the cell death signals of different cell types. Both data types indicated the consistency of SARS-CoV-2 signal and cell death signals, supported by the upregulated cell death-related genes in SARS-CoV-2 RNA-positive cells (Fig. 2e) and the co-localization of SARS-CoV-2 RNAs and cell death-related genes in Stereo-seq data (Fig. 2i). However, ATII, erythrocytes, and club cells were prioritized by Stereo-seq data to be the top three cell types surrounding those cell death signals within 7.5 μm (Fig. 2j), a distance smaller than the diameters of most cells. In contrast, SARS-CoV-2 RNA-positive ATII, erythrocytes, and club cells were only ranked #20, #29, and #26 **(Supplementary Table 5)**, respectively in 76 subpopulations, at the acute damage phase (d2) according to the scRNA-seq data, extremely lower than that of SARS-CoV-2 RNA-positive AM-Cd36 (**Extended Data Fig.2e**). Such discrepancy may be caused by the requirement of scRNA-seq library preparation, *i.e.*, only live cells are eligible for scRNA-seq, and thus “survivor biases” may be introduced. The *in situ* nature of Stereo-seq may capture more cell death-related events and thus is more suitable to investigate SARS-CoV-2-induced damages. While SARS-CoV-2-positive AM-Cd36 macrophages showed upregulation of cell death-related genes and resulted in depletion of AM-Cd36 from hamster lungs two days later after SARS-CoV-2 infection, it appeared that AM-Cd36 macrophages were more resistant to SARS-CoV-2 damages than ATII according to Stereo-seq data (Fig. 2j).

Besides cell death-related genes, *Mki67* was also upregulated in virus-positive AM-Cd36 cells (Fig. 2e), and consequently, we identified a proliferating AM-Cd36 subpopulation termed as AM-Cd36-Mki67 (cluster 19) at d2, the early stage of infection (**Fig.2a-c**). Proliferating AM cells were not observed in healthy controls, and *Mki67* was not observed in other cell populations at d2, enabling us employ this surrogate marker to investigate the spatial distribution of AM-Cd36-Mki67. Spatially, the expression of *Mki67* was dispersed across the centimeter-scale lung sections, with SARS-CoV-2-dense and sparse regions showing similar occurrence (**Extended Data Fig.4a**), suggesting that the AM-Cd36 proliferation may be *in situ* triggered by the viruses as a quick response. Because AMs are the first line of pulmonary defense that survey the lumen of respiratory tracts, our observations suggested that the local host innate immune system may prepare the arms race against SARS-CoV-2 via the *in situ* proliferation of AMs to drive accumulation of lung macrophages through self-renewal, which may occur ahead of the recruitment of potentially tissue-damaging inflammatory cells^22^. However, over-loaded viral particles may break the defense line imposed by AMs via inducing cell deaths of AMs (Fig. 2e-g).

In addition to ATII, erythrocytes, club cells and AM-Cd36 macrophages, lymphatic endothelial cells (LECs), vein endothelial cells (VECs), fibroblast, neutrophils, IMs and T/NKT cells were also responsible for SARS-CoV-2-induced acute tissue damages according to Stereo-seq and scRNA-seq data (Fig. 2j, **Extended Data Fig.4b-d**). SARS-CoV-2-positive cells of these cell types upregulated genes related to cell death (*Bax*, *Bak1*, *Bcl2*, *Daxx* or *Foxo3*), whose cell numbers were decreased at d2 or d5 according to scRNA-seq data, and spatially the marker genes of these cells were enriched in the 7.5 μm neighborhood of spots with cell death signals (**Fig.2j**).

### Acute infection of naïve T cells by SARS-CoV-2 and induction of T cell deaths

Lymphopenia is frequently reported in COVID-19 patients and is predictive of severity^23^. However, it is still unknown how lymphopenia occurs during SARS-CoV-2 infection. In particular, it is unclear whether SARS-CoV-2 infection directly induces lymphocyte deaths for lymphopenia occurring or lymphopenia is only a net result of host immune activation-induced cell deaths. T cell subpopulations demonstrated different temporal changes (**Extended Data Fig.5a**). Although the detection rates of SARS-CoV-2 genes in T cells were relatively lower than those of macrophages and neutrophils, we noticed that the proportion of virus-positive naïve T cells (cluster 59, *Cd3g*^+^*Ccr7*^+^*Tcf7*^+^) in total virus-positive cells was much higher than those of epithelial cell populations at d2 according to our scRNA-seq data (**Extended Data Fig.5b,c**). Similar to the scenarios in epithelial cells, active subgenomic transcription of SARS-CoV-2 was also observed in naïve T cells, with the detection rates of SARS-CoV-2 genes increased along the genome from 5’ to 3’ (**Extended Data Fig.5d**). These findings are consistent with our observations in human bronchial lavage fluid samples based on scRNA-seq, in which SARS-CoV-2 RNAs and subgenomic transcription in T cells were detected^13^. Different from previous studies, our current samples were derived from SARS-CoV-2-infected hamster lungs instead of lavage fluid, and we revealed that naïve T cells instead of activated T cells were the major victims of SARS-CoV-2 infection (**Extended Data Fig.5b,c**), indicating potential escape of SARS-CoV-2 from host adaptive immunity via direct infection and reduction of naïve T cells.

Differential gene expression analysis suggested that *Mx1*, *Isg15* and *Irf7* were upregulated in virus-positive naïve T cells, suggesting cellular intrinsic immune responses to viral infection (Fig. 3a**, Extended Data Fig.5e,f, Supplementary Table 6**). Genes related to cell death, such as *Bax, Daxx, Sod2* and *Lgals1*, were also upregulated in these naïve T cells (Fig. 3a**, Extended Data Fig.5e,f**), indicating the deaths of naïve T cells with SARS-CoV-2 infection. Consistently, the number of naïve T cells significantly decreased at d5 and almost diminished at d7 after SARS-CoV-2 infection (Fig. 3b). These results suggested that the naïve T cells infected with SARS-CoV-2, or at least invaded by SARS-CoV-2, could be induced to cell deaths. In contrast, few virus-positive activated T cells were detected, further indicating such kind of T cell deaths before T cell activation. Together, these phenomena suggest that the lymphopenia observed in COVID-19 patients may be caused, at least partially, by SARS-CoV-2 infection-induced deaths of naïve T cells.

**Fig. 3.**
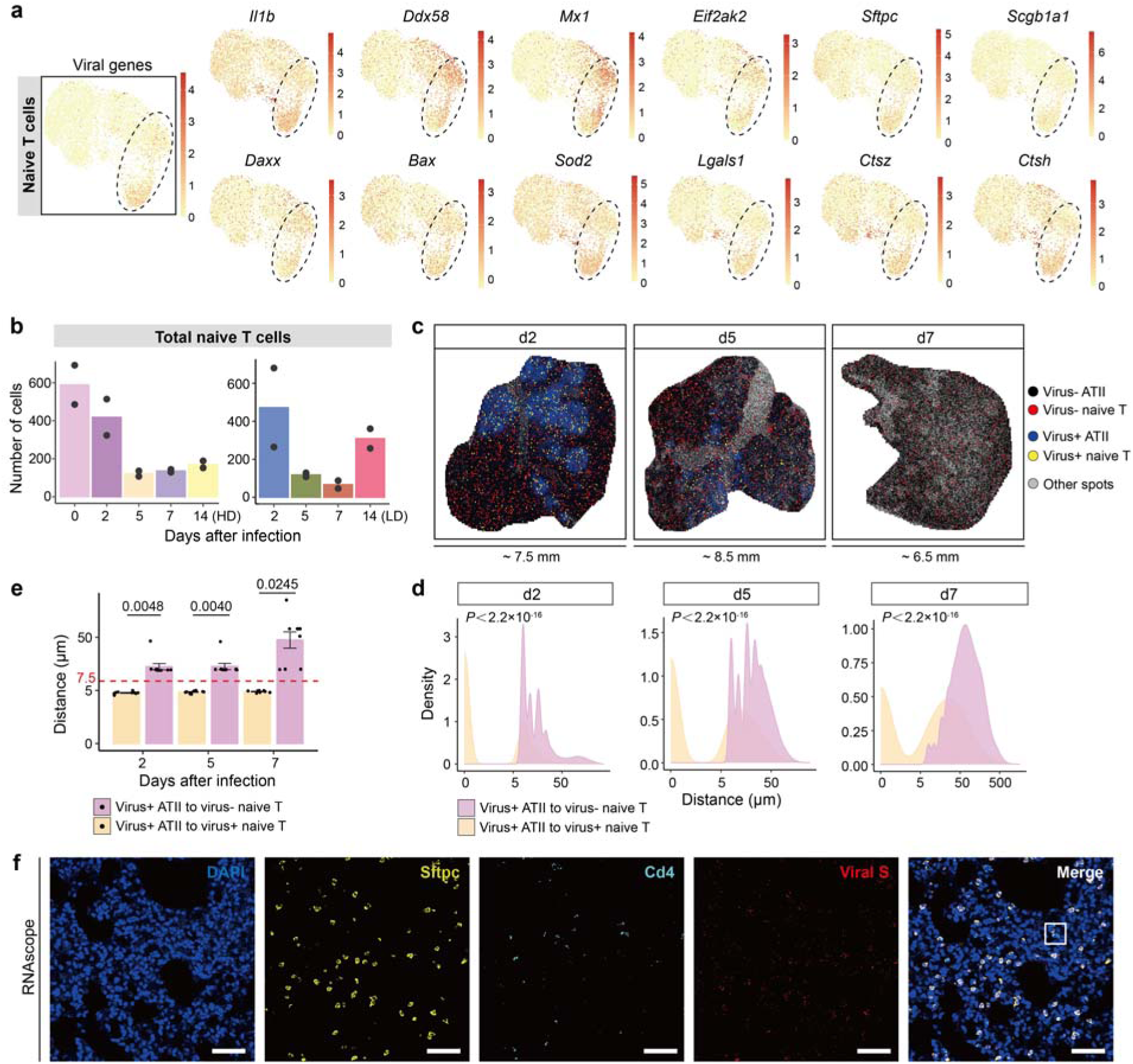
SARS-CoV-2 invades naïve T cells and induces T cell deaths. **a.** The expression of selected differential expressed genes in virus-positive and virus-negative naïve T cells. **b.** Number of total naïve T cells detected at each timepoint. **c.** Spatial distribution of virus-positive/negative naïve T cells around ATII represented by a lung section at d2, d5 and d7 (HD group). The size of virus+ and virus-naïve T spots was magnified for clarity. **d.** Distance between virus-positive ATII cells and their nearest virus-positive/negative naïve T cells in nine sections of d2, d5 and d7 (HD group) respectively. Kolmogorov-Smirnov test. **e.** The peak values of the minimum distance calculated in **(d)** of nine lung sections of each timepoint. Data are mean ± SEM. Two-sided Student t-test. **f.** Staining of virus-positive CD4 T cells and virus-positive ATII by RNAscope. Scale bar, 50 μm.

Is it possible to utilize the spatial transcriptomics data for looking for the way how SARS-CoV-2 infects or invades naïve T cells? As expected, *Ace2* or *Tmprss2*, the host factors proved to be crucial to SARS-CoV-2 entry^21,24^, was not expressed in naïve T cells (**Extended Data Fig.5g**). However, we noticed that *Sftpc* and *Scgb1a1,* marker genes of ATII cells, were upregulated in virus-positive naïve T cells (**Extended Data Fig.5h**). It was intriguing to explore whether cell fusion or exosome-mediated cell-to-cell communication between naïve T cells and ATII contributed to the presence of SARS-CoV-2 RNA-positive naïve T cells. Based on our spatial transcriptomic data with centimeter-scale and subcellular resolution, which enables precise quantification of cell-cell distance together with sufficient statistical power, we examined the spatial relationship between naïve T cells and ATII, and noticed that virus-positive, but not virus-negative naïve T cells, were mostly surrounded by virus-positive ATII (Fig. 3c). The average distance of virus-positive naïve T cells to their nearest virus-positive ATII was shorter than 7.5 μm, in contrast with those of virus-negative naïve T cell at d2, d5 and d7 (Fig. 3d,e). The same analyses were applied to ATI and club cells but ATI cells did not show similar spatial closeness with naïve T cells (**Extended Data Fig.5i,j**). The adjacency of virus-positive naïve T cell and ATII was further validated by *in situ* RNA hybridization (Fig. 3f). As SARS-CoV-2 RNA-positive naïve T cells were also observed by scRNA-seq, the spatial adjacency of virus-positive naïve T cells with ATII within 7.5 μm proposed the possibility that debris or microvesicles generated by ATII after SARS-CoV-2 infection may enter or invade naïve T cells, or SARS-CoV-2-infected ATII may fuse with naïve T cells in the inflamed niche, resulting in SARS-CoV-2 infection of naïve T cells and further causing lymphopenia via T cell deaths or macrophage-mediated phagocytosis.

### Accumulated *Cd38*^+^ and proliferating *Slamf9*^+^*Spp1*^+^ macrophages replenish hamster lung to restrain SARS-CoV-2 infection

While the proportion of lung-resident AM-Cd36 decreased to extremely low levels at d5 and d7 after SARS-CoV-2 infection, two types of monocyte-derived macrophage subpopulations with extremely high detection rates of SARS-CoV-2 genes accumulated in hamster lungs and became the major cell populations of SARS-CoV-2 RNA-positive cells (**Fig.2a-c**). At d2, Mac-Cd38 (cluster 22) and Mac-Slamf9-Spp1 (cluster 27) occupied a small proportion of the myeloid cells within hamster lungs. In particular, only two cells were detected for Mac-Slamf9-Spp1 in HD group by scRNA-seq, but both were SARS-CoV-2 RNA-positive (Fig. 4a). At d5, Mac-Cd38 became the major cell population of SARS-CoV-2 RNA-positive cells instead of AM-Cd36, occupying ∼20% of the entire virus-positive cells (**Extended Data Fig.2e**). The cell surface glycoprotein CD38 has been reported to be an inflammatory marker of human monocytes and macrophages, and to be involved in COVID-19 pathophysiology^25,26^. Accumulation of Mac-Cd38 indicated the severely inflamed status of the lung at d5.

**Fig. 4.**
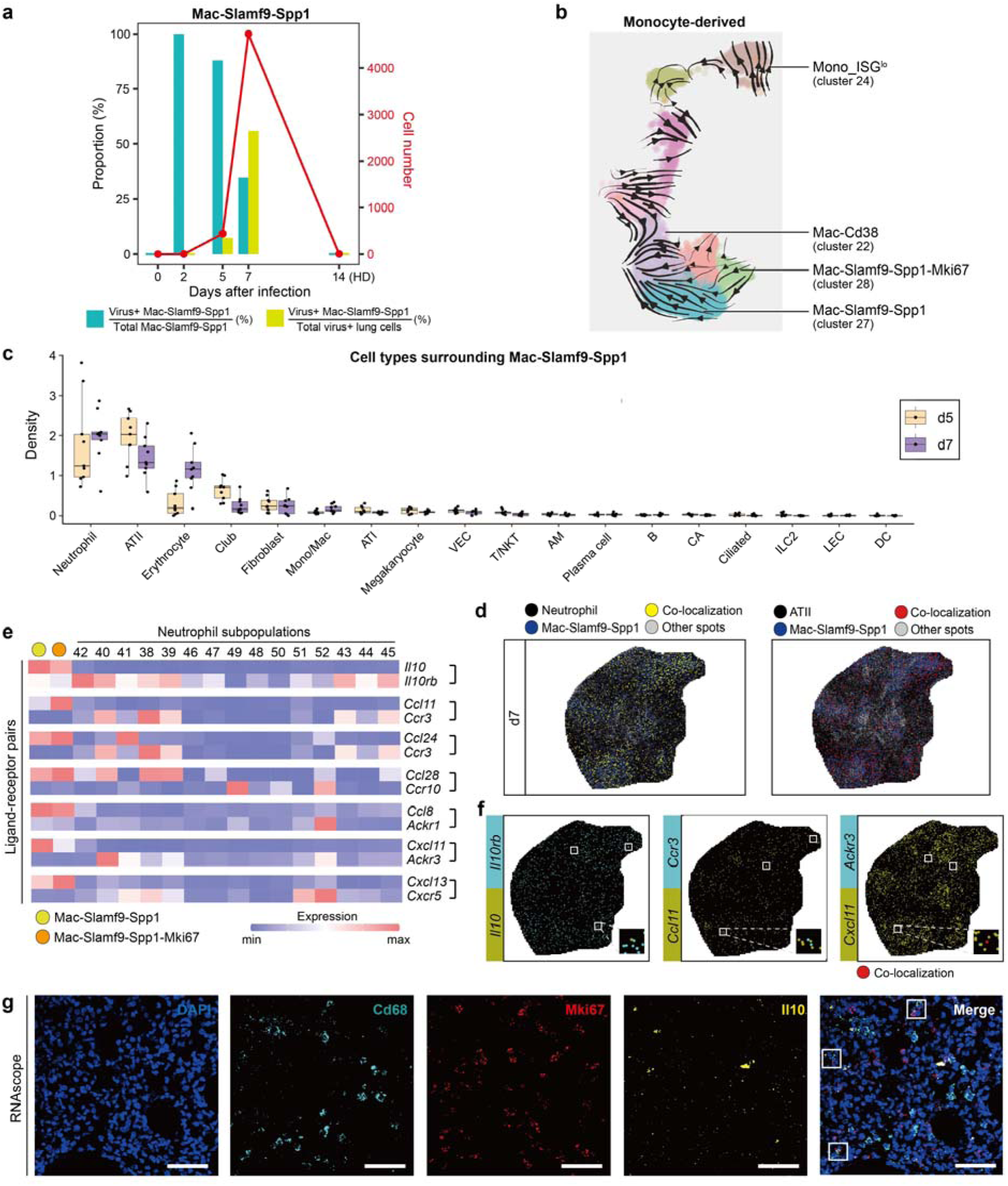
SARS-CoV-2-induced generation of *Slamf9*^+^*Spp1*^+^ macrophages for virus clearance. **a.** The number of total *Slamf9*^+^*Spp1*^+^ macrophages, the proportion of virus-positive *Slamf9*^+^*Spp1*^+^ macrophages and their proportion in total virus-positive cells at each timepoint (HD group). **b.** RNA velocity analysis of mono/macrophage subpopulations. Arrows indicate the potential directions of state transitions. **c.** Density of different cell types in the neighborhood of Mac-Slamf9-Spp1 of nine lung sections at d5 and d7 (HD group). Neighborhood: <7.5 μm. Boxplots: center line, median; box bounds, first and third quartiles; whiskers, 1.5 times the interquartile range. **d.** Spatial distribution of Mac-Slamf9-Spp1, neutrophils and ATII represented by a lung section at d7 (HD group). Yellow spots represented the colocalization of Mac-Slamf9-Spp1 and neutrophils, and red spots represented the co-localization of Mac-Slamf9-Spp1 and ATII. Magnified size of spots was applied for clarity. **e.** Expression of critical ligand and receptor genes in Mac-Slamf9-Spp1, Mac-Slamf9-Spp1-Mki67 and neutrophils subpopulations. Gene description: *LOC101839133*, *Ccl11*; *LOC101824449*, *Ccl8*. **f.** Spatial expression of paired ligand and receptor genes represented by a lung section at d7 (HD group). Co-localization or adjacency of paired genes was framed. **g.** Staining of proliferating *Slamf9+Spp1+* macrophages of lung sections at d7 by RNAscope. Scale bar, 50 μm.

The signaling lymphocytic activation molecule (SLAM) family surface proteins have been proved to act as either inhibitory receptors or activating receptors to regulate immune responses^27,28^. SLAMF9, SLAM family member 9, has been previously identified in human T cells, B cells, NKs and DCs, whose function in immune responses remained mostly unexplored^28^. SPP1, osteopontin, has been reported to play an important role in lung fibrosis^29^. Notably, we identified a new macrophage subpopulation highly expressing *Slamf9, Spp1*, *Siglec1* and *Il10.* These *Slamf9*^+^*Spp1*^+^ macrophages (Mac-Slamf9-Spp1) still maintained an extremely high rate of SARS-CoV-2 RNA positivity at d5 (88%), and the cell number of this cluster increased extremely compared with the two cells detected at d2, occupying 7.3% of all the SARS-CoV-2 RNA-positive cells at d5 (Fig. 4a**, Extended Data Fig. 6a**). Considering the potential false negatives, such a high rate of SARS-CoV-2 RNAs in *Slamf9*^+^*Spp1*^+^ macrophages may suggest that this macrophage subpopulation is specifically induced by engulfing SARS-CoV-2 particles or SARS-CoV-2-infected cells and debris. At d7, *Slamf9*^+^*Spp1*^+^ macrophages further increased and became the major part of SARS-CoV-2 RNA-positive cells, but the SARS-CoV-2 RNA-positive rate of *Slamf9*^+^*Spp1*^+^ macrophages also decreased to 34.8%, occupying 56% of the total virus-positive cells (**Fig.4a, Extended Data Fig. 6a**), suggesting its role in limiting and clearing SARS-CoV-2 infection. The similar tendency was also observed in the LD group (**Extended Data Fig. 6b**). *Cd38*^+^ macrophages and *Slamf9*^+^*Spp1*^+^ macrophages were both characterized by active subgenomic transcription of SARS-CoV-2 (**Extended Data Fig. 6c**). Functional enrichment analysis based on gene ontology revealed that virus-positive *Slamf9*^+^*Spp1*^+^ macrophages primarily upregulated genes related to “tissue remodeling” and “vascular endothelial growth factor production” but not cell death-related genes (**Extended Data Fig. 6d,e, Supplementary Table 6**), implying the cell death-resistant nature of these macrophages and their potential roles in resolving inflammation and viral infection by pro-fibrosis.

Similar to AM-Cd36 cells, proliferation was also observed in *Slamf9*^+^*Spp1*^+^ macrophages (**Fig.2a-c**), confirming the importance of macrophage proliferation in the arms race against viral infection. However, different from virus-positive AM-Cd36 cells, proliferating *Slamf9*^+^*Spp1*^+^ macrophages (Mac-Slamf9-Spp1-Mki67, cluster 28) were not observed to express cell death-related genes, supporting the role of proliferating *Slamf9*^+^*Spp1*^+^ macrophages in limiting and clearing SARS-CoV-2 infection besides its role in resolving inflammation.

We further conducted RNA velocity analysis^30^ based on our scRNA-seq data to interrogate the potential origins and destinations of *Slamf9*^+^*Spp1*^+^ macrophages. The results showed that *Cd38*^+^ macrophages and *Slamf9*^+^*Spp1*^+^ macrophages were derived from peripheral monocytes rather than lung-resident macrophages (**Fig.4b**). Furthermore, mutual state transitions between *Cd38*^+^ macrophages and *Slamf9*^+^*Spp1*^+^ macrophages may exist, probably driven by the engulfment or infection of SARS-CoV-2. Noticing the disappearance of *Slamf9*^+^*Spp1*^+^ macrophages after SARS-CoV-2 clearance at d14 (**Extended Data Fig. 6f**), we speculated that the specific transcriptional state of *Slamf9*^+^*Spp1*^+^ macrophages may be induced by engulfment or infection of SARS-CoV-2, because the viral infection could metabolically and epigenetically reprogram macrophage polarization in certain conditions such as immune evasion and immunosuppression^31,32^. Whether such reprogramming exists and what roles it plays in SARS-CoV-2 infection and clearance needs to be further investigated.

To identify the spatial context of *Slamf9*^+^*Spp1*^+^ macrophages and investigate the potential mechanisms underlying *Slamf9*^+^*Spp1*^+^ macrophages-mediated SARS-CoV-2 clearance, we calculated the density of diverse cell populations surrounding *Slamf9*^+^*Spp1*^+^ macrophages marker genes based on our spatial transcriptomic data at the tissue scale. We found that the density of neutrophils spatially surrounding *Slamf9*^+^*Spp1*^+^ macrophages increased from d5 to d7 and neutrophils formed the nearest cell type within the 7.5 μm neighborhood of *Slamf9*^+^*Spp1*^+^ macrophages at d7, much closer than ATII and erythrocytes (**Fig.4c,d**), suggesting that recruitment of neutrophils and coordination may be important to contain and eliminate SARS-CoV-2 infection. Ligand-receptor analysis based on scRNA-seq data suggested that *Slamf9*^+^*Spp1*^+^ macrophages may have distinct intercellular interactions with different neutrophil subpopulations (**Extended Data Fig. 6g, Supplementary Table 7**). In particular, *Slamf9*^+^*Spp1*^+^ macrophages expressed high levels of *Ccl8*, *Ccl28*, *Ccl11* and *Cxcl11*, which might recruit *Ackr1*^+^ and *Ccr10*^+^ neutrophil precursors from the blood, as well as *Ccr3*^+^ and *Ackr3*^+^ mature neutrophils to their surroundings (**Fig.4e**). Specifically, *Slamf9*^+^*Spp1*^+^ macrophages may interact via the axis of *Il10* and its receptor with acNeu-Isg12-Cst7 (**Fig.4e**), which highly expressed *Il10rb* and was also associated with SARS-CoV-2 clearance (See the next section for details). Spatially, these specific ligand-receptor pairs were co-localized or adjacently distributed according to our Stereo-seq data (**Fig.4f**), supporting the scRNA-seq data observations and suggesting the potential coordination of the different innate cell populations in clearing viruses through these ligand-receptor pairs.

To further verify the presence of the proliferating *Slamf9*^+^*Spp1*^+^ macrophages, we carried out *in situ* RNA hybridization, and observed the co-localization of *Cd68, Il10* and *Mki67* RNA (**Fig.4g**). The result emphasized the possibility of the *in situ* proliferation of lung macrophages to fight against virus or contribute to viral clearance during SARS-CoV-2 infection.

### *Isg12*^+^*Cst7*^+^ neutrophils specifically accumulated for SARS-CoV-2 elimination

During the acute phase of inflammation, neutrophils are generally concerned to be recruited from peripheral blood to inflamed sites to clear infection or tissue injuries^33^. Consistently, we observed that the hamster lungs were replenished with peripherally derived neutrophils and demonstrated high phenotype diversity. A total of 15 subpopulations was identified for neutrophils based on our scRNA-seq data, including precursor, immature, mature, activated neutrophils, and two subpopulations highly expressing *Vcan* and *Lyz* (**Extended Data Fig.7a,b**). Similar to macrophage subsets, these neutrophil subpopulations with different cellular and functional states also demonstrated distinct temporal patterns (**Extended Data Fig.7c,d, Supplementary Table 8**).

Similar to AM-Cd36, lowly-activated neutrophils (cluster 38, acNeu_lo) diminished after infection (**Extended Data Fig.7d**). Highly-activated neutrophils (cluster 40, acNeu_hi) were the major population of SARS-CoV-2 RNA-positive neutrophils at d2 (**Extended Data Fig.7e**). SARS-CoV-2 RNA-positive highly-activated neutrophils upregulated genes related to IL-6 production, and type I IFN production (**Fig.5a, Supplementary Table 6**), indicating the pro-inflammatory role of these neutrophils and potentially contributing to the acute super-inflammation caused by SARS-CoV-2 infection. Genes related to cell death were also observed to be upregulated in virus-positive highly-activated neutrophils (**Fig.5a**). Consequently, the number of such acNeu_hi neutrophils decreased at d5 and d7 (**Extended Data Fig.7f**), suggesting that SARS-CoV-2 infection also damages lung neutrophils besides epithelial cells and lung-resident macrophages.

Clusters 39 and 41 were also enriched at d2 and upregulated *Tnf* and *Cdkn1a* expression (**Extended Data Fig.7b**), suggesting an increased activation state of lung neutrophils. Sequential to lung neutrophil activation and death, a large number of immature neutrophils were chemoattracted and accumulated in the hamster lung tissues at d5 and d7 (**Extended Data Fig.7d**), empowering the host with increasingly augmented innate immune responses.

We noticed that a new subpopulation of neutrophils characterized by high expression of *Isg12, Cst7,Cd74, Slamf7 and Cd274* (acNeu-Isg12-Cst7, cluster 42) was enriched at d7 and formed the major part of total virus-positive neutrophils (Fig. 5b). CST7 (cystatin F), a cysteine peptidase inhibitor, is known to suppress cytotoxicity of T cells and NK cells^34^. ISG12 (interferon-alpha inducible protein 27) is reported to regulate innate inflammatory responses and also inhibit viral replication^35,36^. A negative correlation was observed between the proportion of virus-positive total cells and that of *Isg12*^+^*Cst7*^+^ neutrophils (Fig. 5c), and virus-positive *Isg12*^+^*Cst7*^+^ neutrophils upregulated genes related to vesicle organization and vesicle-mediated transport (Fig. 5a), implying the possibility that these neutrophils could eliminate virus through the transport of extracellular vesicles^37^. Based on the spatial transcriptomic data, we found that, instead of activated neutrophils which located nearby virus, *Isg12*^+^*Cst7*^+^ neutrophils were spatially distant from viral spots (**Fig.5d,e, Extended Data Fig.7g-j**), further confirming the potential function of *Isg12*^+^*Cst7*^+^ neutrophils to eliminate the surrounding viruses or virus-infected cells. Moreover, *Isg12*^+^*Cst7*^+^ neutrophils expressed *Cd274* and *Il10rb* (**Fig.5b**), potentially interacted with activated cytotoxic T cells and *Slamf9*^+^*Spp1*^+^ macrophages and jointly established the favoring niches for the resolution of inflammation (**Fig.4e**).

**Fig. 5.**
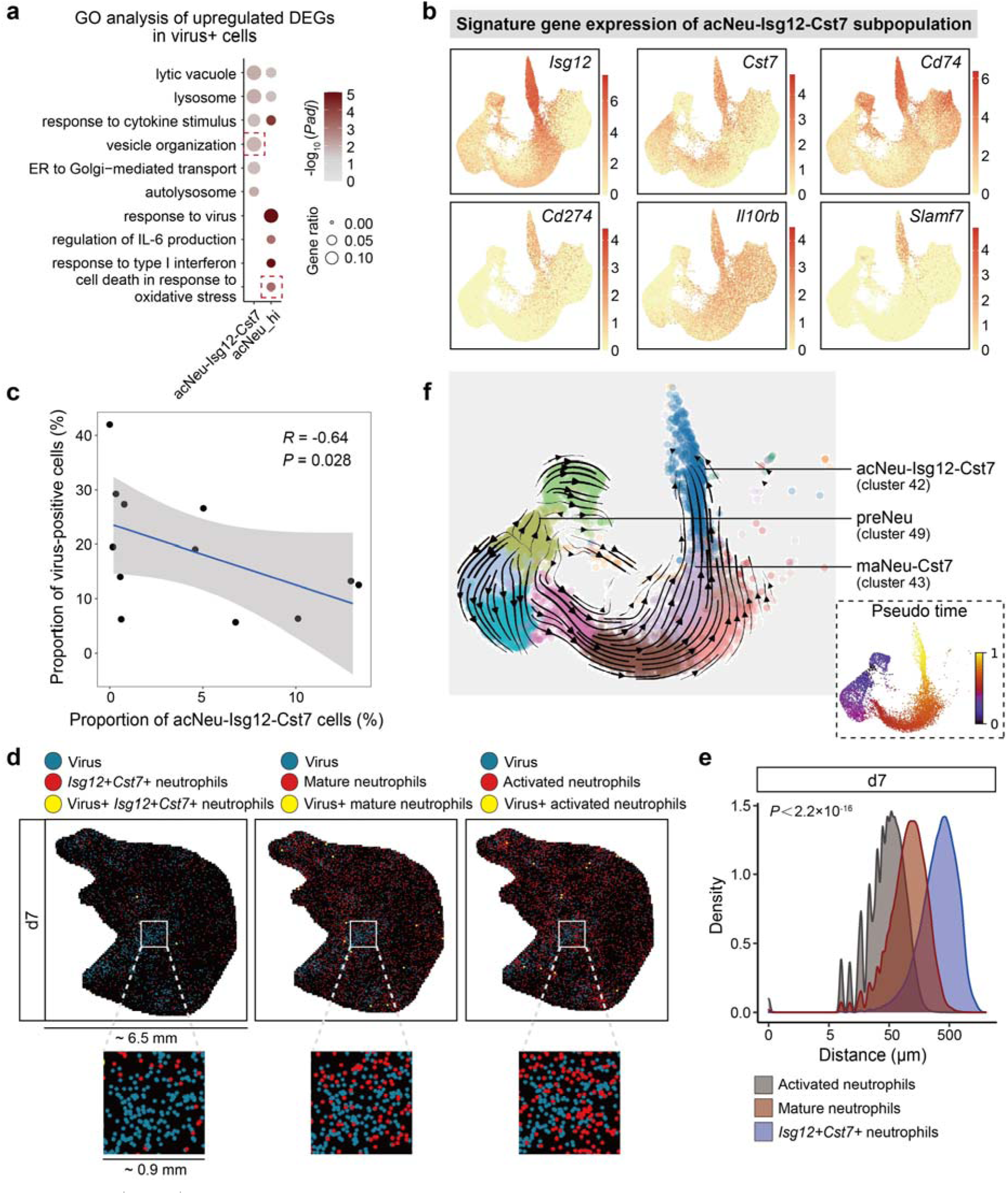
*Isg12+Cst7+* neutrophils accumulate for eliminating virus. **a.** GO terms enriched in upregulated genes of virus-positive highly-activated neutrophils and *Isg12^+^Cst7^+^* neutrophils. **b.** The expression of signature genes identifying acNeu-Isg12-Cst7 subpopulation. Gene description: *LOC101838467*, *Isg12*. **c.** Association between the proportion of virus-positive cells and *Isg12^+^Cst7^+^* neutrophils in 12 hamsters (d2, d5 and d7, HD and LD). Spearman’s correlation. **d.** Spatial distribution of virus and activated, non-activated mature or *Isg12^+^Cst7^+^* neutrophils, represented by a lung section at d7 (HD group). Magnified size of all spots was applied for clarity. **e.** Distance of viral spots to their nearest neutrophil subpopulations of nine sections of d7 (HD group). Kolmogorov-Smirnov test. **f.** RNA velocity analysis of neutrophil subpopulations. Arrow direction indicates the potential directions of state transitions.

To further evaluate the potential origins and destinations of these *Isg12*^+^*Cst7*^+^ neutrophils, we conducted RNA velocity analysis for the neutrophil subpopulations. Similar to those *Slamf9*^+^*Spp1*^+^ macrophages which were peripheral monocytes-derived, we found that *Isg12*^+^*Cst7*^+^ neutrophils were also derived from those neutrophil precursors recruited from peripheral blood (**Fig.5f**). Different from *Slamf9*^+^*Spp1*^+^ macrophages, proliferation was not observed in *Isg12*^+^*Cst7*^+^ neutrophils. Instead, we identified three proliferating subpopulations in neutrophil precursors and immature neutrophils (**Extended Data Fig.7a,b**). RNA velocity analysis suggested that developmental linkages existed among these proliferating neutrophils, non-proliferating immature neutrophils, mature neutrophils, activated neutrophils and *Isg12*^+^*Cst7*^+^ neutrophils (**Fig.5f**). These data suggested that the proliferation may still be an important strategy for the host neutrophils to win the arms race against virus infection but was mainly contributed by the immature populations, contrasting with the proliferation of *Slamf9*^+^*Spp1*^+^ macrophages and AM-Cd36 aforementioned.

### *Trem2*^+^ and *Fbp1*^+^ macrophages enriched for inflammation resolution at the late stage

Both scRNA-seq and Stereo-seq data revealed that SARS-CoV-2 were nearly absent from hamster lungs at d14, suggesting viral clearance by the host at the late stage of infection. Similarly, immune infiltration and lung injury were also attenuated at d14 revealed by pathological observations (**Fig.1c**). Accompanying with the diminishment of SARS-CoV-2, the *Cd38*^+^ macrophages, *Slamf9*^+^*Spp1*^+^ macrophages, and *Isg12^+^Cst7^+^* neutrophils also diminished, which was consistent with their roles in viral clearance. Since *Cd38*^+^ and *Slamf9*^+^*Spp1*^+^ macrophages did not exhibit expression of cell death-related genes, a critical question emerges, *i.e.*, what is the destination of *Cd38*^+^ and *Slamf9*^+^*Spp1*^+^ macrophages after viral clearance. We interrogated the phenotypic similarity between d7 and d14 myeloid cells by mutually mapping cells at one day (Query) to those at the other day (Reference) based on supervised machine learning algorithms implemented in Seurat. The results suggested that tight linkage exists between *Slamf9*^+^*Spp1*^+^ macrophages and *F13a1*^+^, *Trem2*^+^ and *Fbp1*^+^ macrophages (Fig. 6a), indicating that *Slamf9*^+^*Spp1*^+^ macrophages may differentiate into these macrophage subpopulations to promote inflammation resolution and tissue repair.

**Fig. 6.**
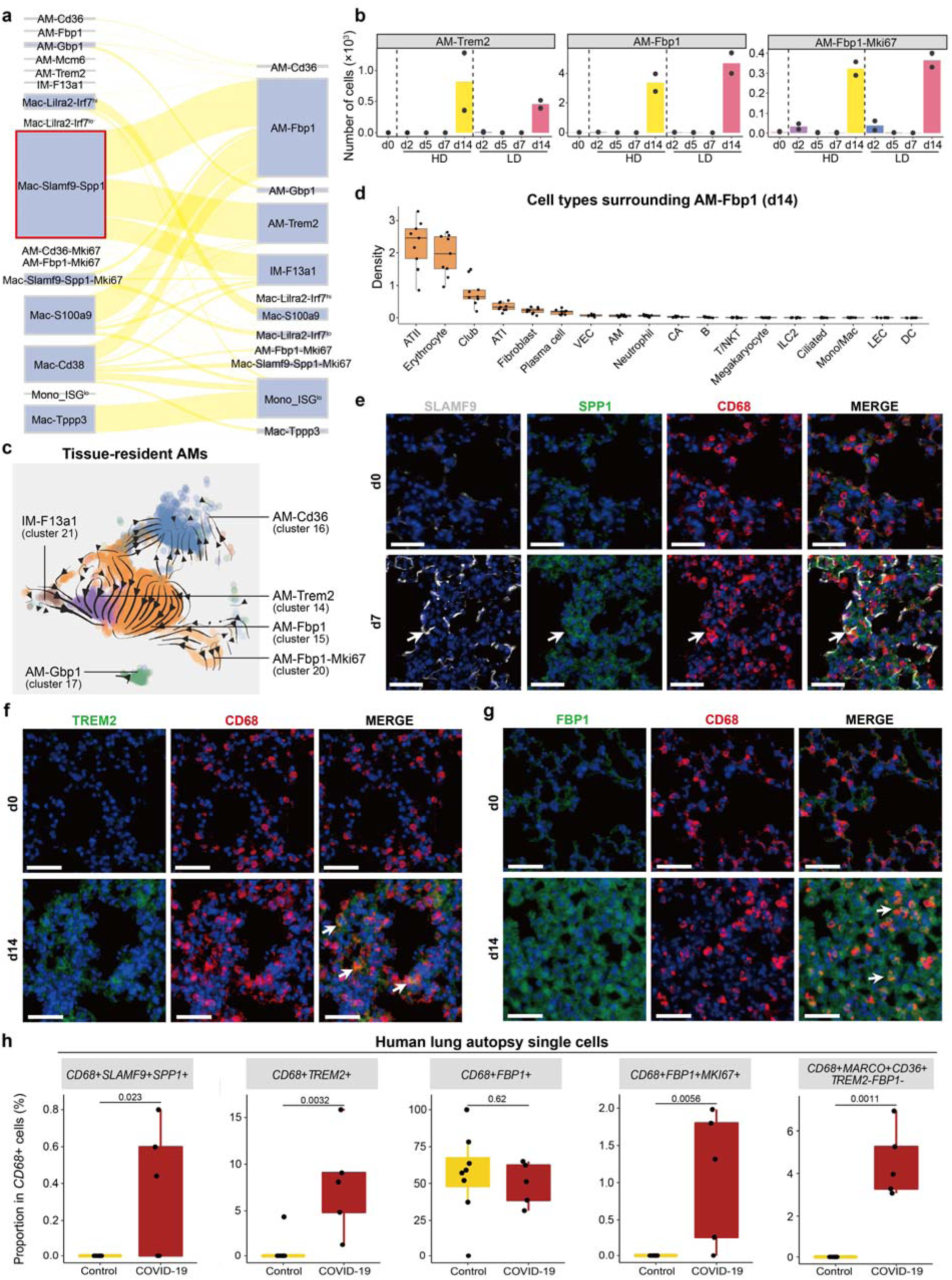
Identification of *Trem2*^+^ and *Fbp1*^+^ alveolar macrophages enriched for inflammation resolution. **a.** Phenotypic linkage between AM and mono/macrophage subpopulations from d7 and d14. **b.** Temporal changes of AM-Trem2, AM-Fbp1 and AM-Fbp1-Mki67. **c.** RNA velocity analysis of AM subpopulations. Arrows indicate the potential directions of state transitions. **d.** Density of different cell types in the neighborhood of AM-Fbp1 of nine lung sections at d14 (HD group). Neighborhood: <7.5 μm. Boxplots: center line, median; box bounds, first and third quartiles; whiskers, 1.5 times the interquartile range. **e-g.** Immunofluorescence analyses of the lung sections of hACE2 mice before and after SARS-CoV-2 infection. Staining of SLAMF9, SPP1 and CD68 at d0 and d7 (**e**); TREM2 and CD68 at d0 and d14 (**f**); FBP1 and CD68 at d0 and d14 (**g**). Scale bar, 50 μm. **h.** Identification of *SLAMF9*^+^*SPP1*^+^, *TREM2*^+^, *FBP1*^+^, *FBP1*^+^*MKI67*^+^, and *MARCO*^+^*CD36*^+^*TREM2*^-^*FBP1*^-^ macrophages in scRNA-seq data of human lung autopsy samples of COVID-19 patients. Control: pulmonary bulla or lung cancer, and these samples were to be inflammation-free pathologically confirmed. Two-sided Wilcoxon test.

Interestingly, *Trem2*^+^ and *Fbp1*^+^ macrophages were specifically enriched at d14 but physiologically absent in healthy controls, which were termed as AM-Trem2 (cluster 14, *Trem2*^hi^ *Spp1*^hi^), AM-Fbp1 and AM-Fbp1-Mki67 (cluster 15 and 20, *Fbp1*^hi^; **Fig.2a-c, 6b**). Different from *Trem2*^+^ and *Fbp1*^+^ macrophages, *F13a1*^+^ macrophages (IM-F13a1, cluster 21) were enriched at d0 and d2 but depleted at d5 and d7 and partially restored at d14 (**Fig.2c**). The expression of *Trem2* (triggering receptor expressed on myeloid cells 2, related to fibrosis^38^) and *Fbp1* (fructose-bisphosphatase 1, a key enzyme in gluconeogenesis) may modulate the metabolic state and the function of macrophages for inflammation resolution and tissue repair after SARS-CoV-2 infection^38,39^.

RNA velocity analysis suggested that sequential state transitions from AM-Trem2 towards AM-Fbp1 and AM-Cd36 may exist (**Fig.6c**). Similar to IM-F13a1, AM-Cd36 macrophages were also enriched at d0 and d2 but depleted from d5 and d7 and partially restored at d14. Mutual mapping between d7 and d14 together with RNA velocity results suggested that further differentiation or polarization of *Slamf9*^+^*Spp1*^+^ macrophages into *F13a1*^+^, *Trem2*^+^ and *Fbp1*^+^ macrophages may be pivotal to inflammation resolution and tissue repair after clearing SARS-CoV-2.

We validated the RNA velocity analysis results by conducting trajectory and pseudo-time analysis based on Monocle2. The results confirmed insights generated by RNA velocity analysis and further indicated genes undergoing prominent temporal changes during the differentiation, e.g., *Trem2*, *Spp1*, *C1qa*, *Fcgr2b*, *Tnf*, *Cxcl14*, *Marco*, *Cd36*, etc (**Extended Data Fig.8a-c**), consistent with the process from inflammation resolution and tissue repair.

Evidenced by the spatial transcriptomics data, ATII and erythrocytes were the top two cell types enriched in the neighborhood of the new subpopulation of *Fbp1*^+^ AMs (**Fig.6d**), indicating the roles in tissue repair. Notably, plasma cells were observed to be relatively enriched around *Fbp1*^+^ AMs (**Fig.6d**). These plasma cells expressed *Odc1*, *Btla* and *Slamf7* (**Extended Data Fig.8d**), indicating their potential roles in restraining inflammation. These plasma cells also upregulated *Prdx4*, *Cd74* and *Apoe*, compared to those at d0 (**Extended Data Fig.8e**), suggesting their immunosuppressive functions.

The identification of proliferating AM-Fbp1 new subpopulation (AM-Fbp1-Mki67, cluster 20) at d14 may again suggest the importance of macrophage proliferation in inflammation resolution and tissue repair in addition to the arms race against viral infection^22^.

We validated the presence of these *Fbp1*^+^ AMs and *Trem2*^+^ AMs specifically emerging at the inflammation resolution stage (d14) by *in situ* RNA hybridization (**Extended Data Fig.8f**). The existence of *Slamf9*^+^*Spp1*^+^, *Trem2*^+^ and *Fbp1*^+^ macrophages identified in our data were further supported by immunofluorescence analyses in the lung sections of authentic SARS-CoV-2-infected, human ACE2 transgenic mice (hACE2 mice) **Fig.6e-g**). Consistent with our observations in hamsters, SLAMF9^+^SPP1^+^CD68^+^ cells were observed at d7 and TREM2^+^CD68^+^ and FBP1^+^CD68^+^ cells at d14 in hACE2 mice after SARS-CoV-2 infection, while all these cells were not found at d0. In particular, we validated the presence of these macrophages based on a publically available human autopsy scRNA-seq dataset^40^. By employing a set of signature genes specifically expressed in *Slamf9*^+^*Spp1*^+^, *Trem2*^+^, *Fbp1*^+^, and *Marco*^+^*Cd36*^+^ macrophages identified in our study (**Extended Data Fig.8g**), we confirmed the enrichment of *SLAMF9*^+^*SPP1*^+^, *TREM2*^+^, *FBP1*^+^, and *MARCO*^+^*CD36*^+^ macrophages and even the proliferation of *FBP1*^+^ macrophages in lung autopsy samples of COVID-19 patients (**Fig.6h**). In contrast, autopsy samples from pulmonary bulla or lung cancers that were pathologically confirmed to be inflammation-free demonstrated only *FBP1*^+^ macrophages. These results suggest that the roles of *Slamf9*^+^*Spp1*^+^ macrophages in SARS-CoV-2 clearance, inflammation resolution, and tissue repair observed in hamsters may be translated to humans and thus are significant for developing effective COVID-19 therapeutics to control the current pandemic.

## Discussion

Taken together, our spatiotemporal study of SARS-CoV-2 pulmonary infection revealed a new subpopulation of macrophages, *i.e.*, *Slamf9*^+^*Spp1*^+^ macrophages, which is at the core of SARS-CoV-2 clearance and thereafter inflammation resolution and tissue repair. The dynamics and mechanisms of *Slamf9*^+^*Spp1*^+^ macrophages in clearing SARS-CoV-2 and inflammation may be illustrated as **Extended Data Fig. 9** and summarized as follows: 1) derived from peripheral monocytes with Mac-Cd38 as intermediates; 2) induced specifically by SARS-CoV-2 engulfment or infection because of the extremely high rates of SARS-CoV-2 RNA positivity in these cells; 3) actively proliferating to win the arms race against SARS-CoV-2 intrahost transmission; 4) cell death-resistant, different from lung-resident alveolar macrophages; 5) secreting chemokines Ccl11 and Cxcl11 to recruit neutrophils and clear SARS-CoV-2 together; 6) secreting IL-10 to induce an immunosuppressive niche that is co-localized by *Isg12*^+^*Cst7*^+^ neutrophils to prevent inflammation-induced damages; 7) differentiating into *Trem2*^+^ and *Fbp1*^+^ macrophages to further resolve inflammation; and 8) replenishing IMs directly or AMs indirectly.

While our study is limited by the observational nature based on Stereo-seq and scRNA-seq data and future studies are needed to confirm and substantiate these observations, it is worth noting that such findings were directly derived from *in vivo* SARS-CoV-2 infection with the lens of Stereo-seq and scRNA-seq. When a new infectious disease emerged, it is of urgent need to quickly identify those host factors that are important to limit, clear the pathogens and resolve their damages. However, because of the complexity of in vivo infection and the related host immune responses, traditional technologies cannot address such challenges due to the limitations of spatial, temporal, and cellular resolution, limiting the development of anti-infectious measurements and therapies. Although scRNA-seq provides sufficient resolution at the single-cell level, the loss of spatiotemporal information and the “survivor biases” introduced during scRNA-seq library preparation as we illustrated in this study prohibit its power to directly identify critical host factors important for pathogen clearance and inflammation resolution. Our study demonstrated the power of spatial omics technologies in illustrating the molecular and cellular details of in vivo biological events, specifically those locally related to cell damages and cell deaths. Therefore, single-cell spatiotemporal studies of in vivo biological phenomena will be a powerful paradigm to revolutionize studies including inflammatory damages, infectious diseases, tumours, and any other questions related to cell deaths.

As single-cell spatiotemporal studies are inherently unbiased and observational, the generated hypotheses will be more reliable than guesses based on limited indices and more deserving to be validated and deepened by experimental studies. Our study revealed that *Slamf9*^+^*Spp1*^+^ macrophages were specifically induced by SARS-CoV-2 infection. However, the conditions sufficient or necessary for the induction of *Slamf9*^+^*Spp1*^+^ macrophages are still unknown and further studies are needed to illustrate the metabolic, epigenomic, physical factors necessary for the induction and function of *Slamf9*^+^*Spp1*^+^ macrophages. Similarly, proliferation of different macrophage subsets and neutrophils has been observed to be an important mechanism to act against SARS-CoV-2 infection.

However, the factors and conditions related to the macrophage and neutrophil proliferation in vivo are still unknown and require future experiments to illustrate. While the identification of naïve T cells undergoing SARS-CoV-2 invasion or infection by incorporating SARS-CoV-2-infected ATII-derived debris or microvesicles^41^ has been insightful based on the observations provided by Stereo-seq and scRNA-seq data, the detailed mechanisms and the relationship with clinical lymphopenia are still important questions requiring future experimental studies to address. *Cd274*^+^ and *Cst7*^+^ neutrophils were previously observed in the blood sample of COVID-19 patients and associated with disease severity^13,42^, confirming the clinical relevance of our finding and indicating the importance of this group of neutrophils in understanding COVID-19. But the detailed mechanisms of this group of neutrophils in SARS-CoV-2 clearance still need further experiments to substantiate. Since long COVID-19 has increasingly become a global concern^43^, the identification of *Trem2*^+^ and *Fbp1*^+^ macrophages by our study and the illumination of their roles in inflammation resolution and tissue repair is helpful to develop treatment of post-acute infection sequelae of COVID-19 but the underlying factors critical for the functions of these macrophage subgroups are still pending to clarify.

In summary, our study depicted the spatiotemporal changes of SARS-CoV-2 pulmonary infection in hamsters at single-cell resolution and characterized the cellular and molecular events covering the whole process from viral infection, tissue damage, immune infiltration, viral clearance, to inflammation resolution and tissue repair. The identification of *Slamf9*^+^*Spp1*^+^ macrophages and their potential descendant cells together with *Isg12*^+^*Cst7*^+^ neutrophils, which were derived from infiltrated peripheral immune precursors, provided actionable hypotheses for precisely understanding the mechanisms underlying viral clearance and inflammation resolution, which is insightful for developing anti-virus therapies. In particular, the identification of virus-positive naïve T cells and their fates towards cell deaths as well as their spatial co-localization with virus-positive ATII cells shed light on lymphopenia that is associated with COVID-19 severity in humans. The rich information of our scRNA-seq and Stereo-seq data provides a comprehensive and insightful resource for understanding viral diseases and developing anti-viral therapies in future.

## Data availability

The data that support the findings of this study have been deposited into CNGB Sequence Archive (CNSA) of China National GeneBank DataBase (CNGBdb) with accession number CNP0002742 and CNP0002978. All other data are included in the article and/or supporting information.

## Acknowledgements

We thank Young Li, Yijian Li, Xiaoyu Wei, Hongwei Pan, Chao Chen, Ninghe Wang, Yumo Song for technical assistant. This work was supported by Grants from the National Natural Science Foundation of China (81788101 and 32022016), CAMS Innovation Fund for Medical Sciences (2021-I2M-1-017), National Key R&D Program of China (2018YFA0507400) and Emergency Key Program of Guangzhou Laboratory (EKPG21-30-3).

## Author contributions

X.C., J.W., L.L., C.Q. designed the experimental approach and supervised the study. B.C., P.Y., W.D. and F.L. performed animal experiments. B.C., X.D., J.K., Y.C., Q.Y., J.L., Y.F., and Y.L. carried out the library preparation and other relevant experiments. B.C., J.L. and Y.G. performed RNAscope experiments. Z.L. and S.L. provided the reagents. Z.Y., Y.Z., Y.F., and Y.L. performed data preprocessing and quality control. Y.C., J.L. and M.Z. helped analyze the data. N.Z., W.F. and X.X. provided technical support and helpful discussion. B.C, Z.Y., L.L., X.R. and X.C. analyzed the data and wrote the paper with all authors’ inputs.

---

**Extended Data Fig.1.**
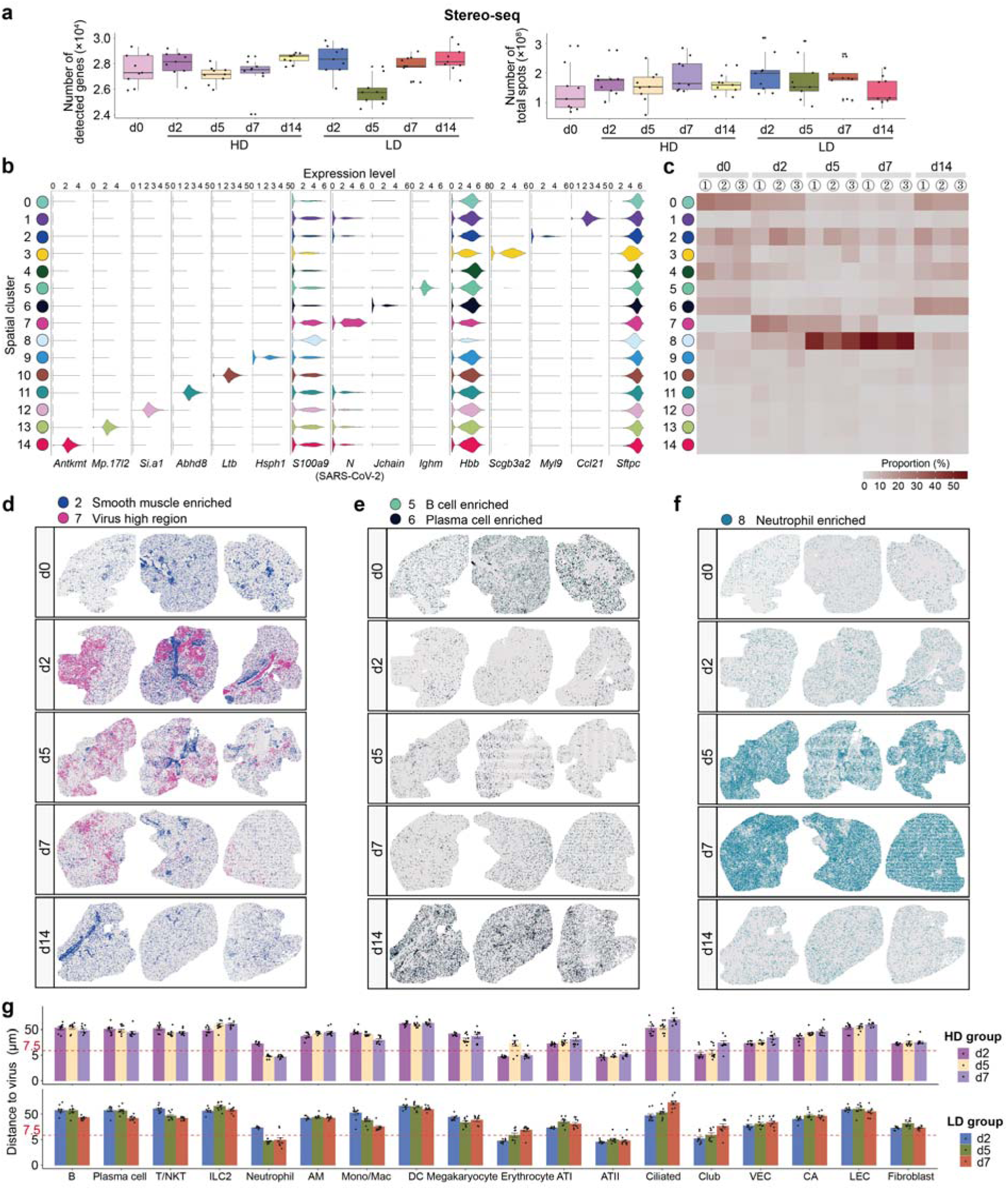
Spatial transcriptomics of hamster lung sections. **a.** Quality control of Stereo-seq data. Total genes detected and total spot numbers of 81 spatial transcriptomic sections (9 sections per timepoint). **b.** The expression of selected marker genes to annotate different spatial regions in each spatial cluster. **c.** The proportion of each spatial cluster in total spots of each section. **d-f.** The highlight of specific spatial clusters. Smooth muscle enriched region and virus high region (**d**); B cell enriched and plasma cell enriched region (**e**); neutrophil enriched region (**f**). **g.** The peak values of the minimum distance between virus and the nearest cell types, calculated in the nine lung sections of each timepoint.

**Extended Data Fig.2.**
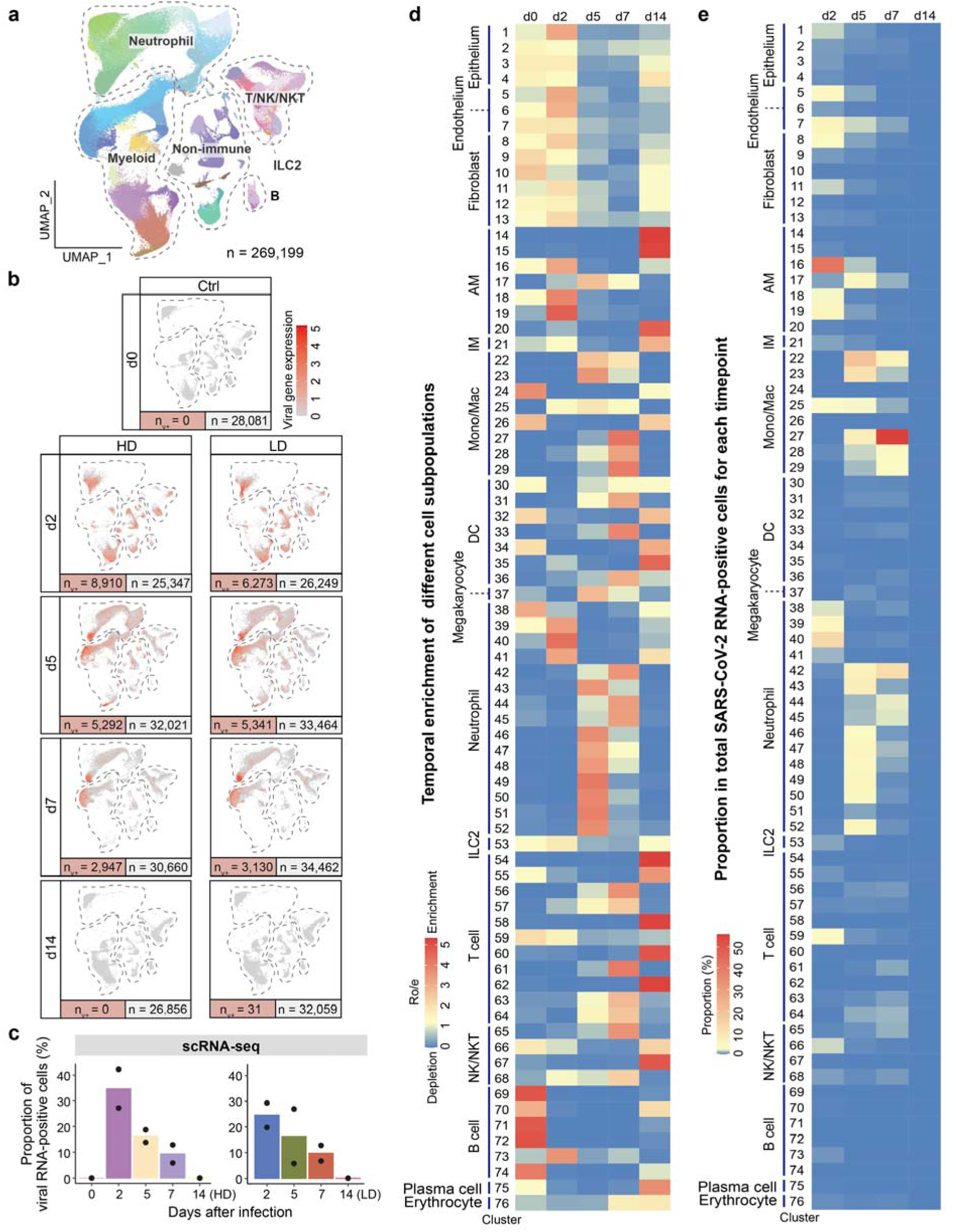
A single-cell landscape of SARS-CoV-2-infected hamster lungs identifying 76 subpopulations. **a.** UMAP projection of hamster lung cells (n=269,211) showing 76 cell subpopulations. Non-immune (epithelium, endothelium, fibroblast, erythrocyte); Myeloid; Neutrophil; ILC2; T/NK/NKT; B. **b.** Distribution of SARS-CoV-2 RNA-positive cells (n, number of total cells; n_v+_, number of virus-positive cells). **c.** Proportion of SARS-CoV-2 RNA-positive cells (viral count>0) at each timepoint. **d.** Temporal distributions of all 76 scRNA-seq cell subpopulations at d0, d2, d5, d7 and d14 (HD group). Ro/e > 1, enrichment; Ro/e < 1, depletion. **e.** Distribution of SARS-CoV-2 RNA-positive cells across 76 cell subpopulations at d2, d5, d7 and d14 (HD group).

**Extended Data Fig.3.**
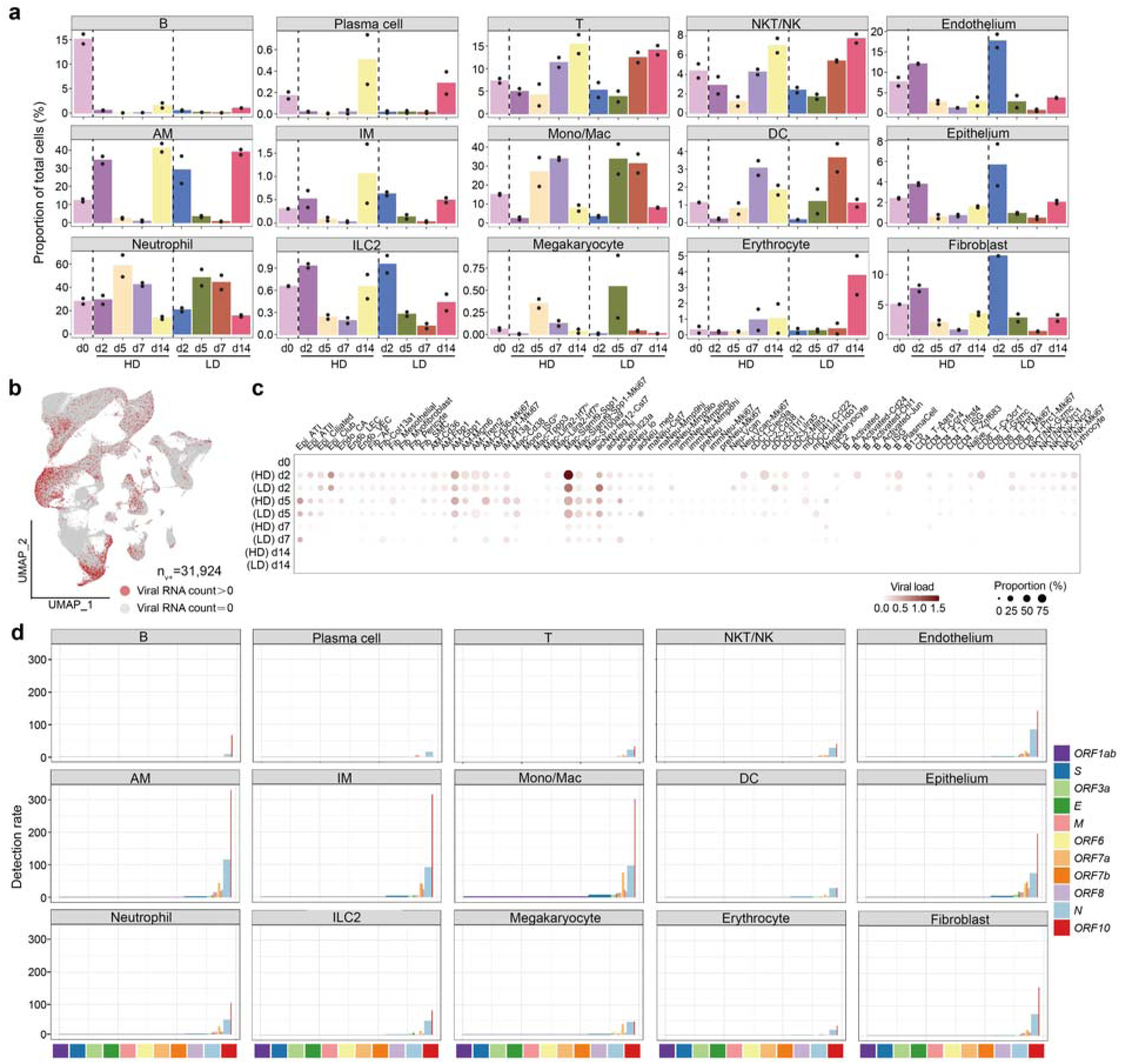
Signatures of SARS-CoV-2 RNA-positive cells. **a.** The proportion of various cell types of two replicated hamsters at each timepoint. **b.** Distribution of 31,924 SARS-CoV-2 RNA-positive cells (n_v+_, number of virus-positive cells). **c.** The relative proportion of virus-positive cells within 76 cell subpopulations. **d.** Detection rates of SARS-CoV-2 genes in main cell types. Given a viral gene *g_v_*, the detection rate is defined as the ratio of the number of *g_v_*^+^ cells to the total cells of the specific cell type and then normalized by the gene length in the SARS-CoV-2 genome and multiplied by 10^6^.

**Extended Data Fig.4.**
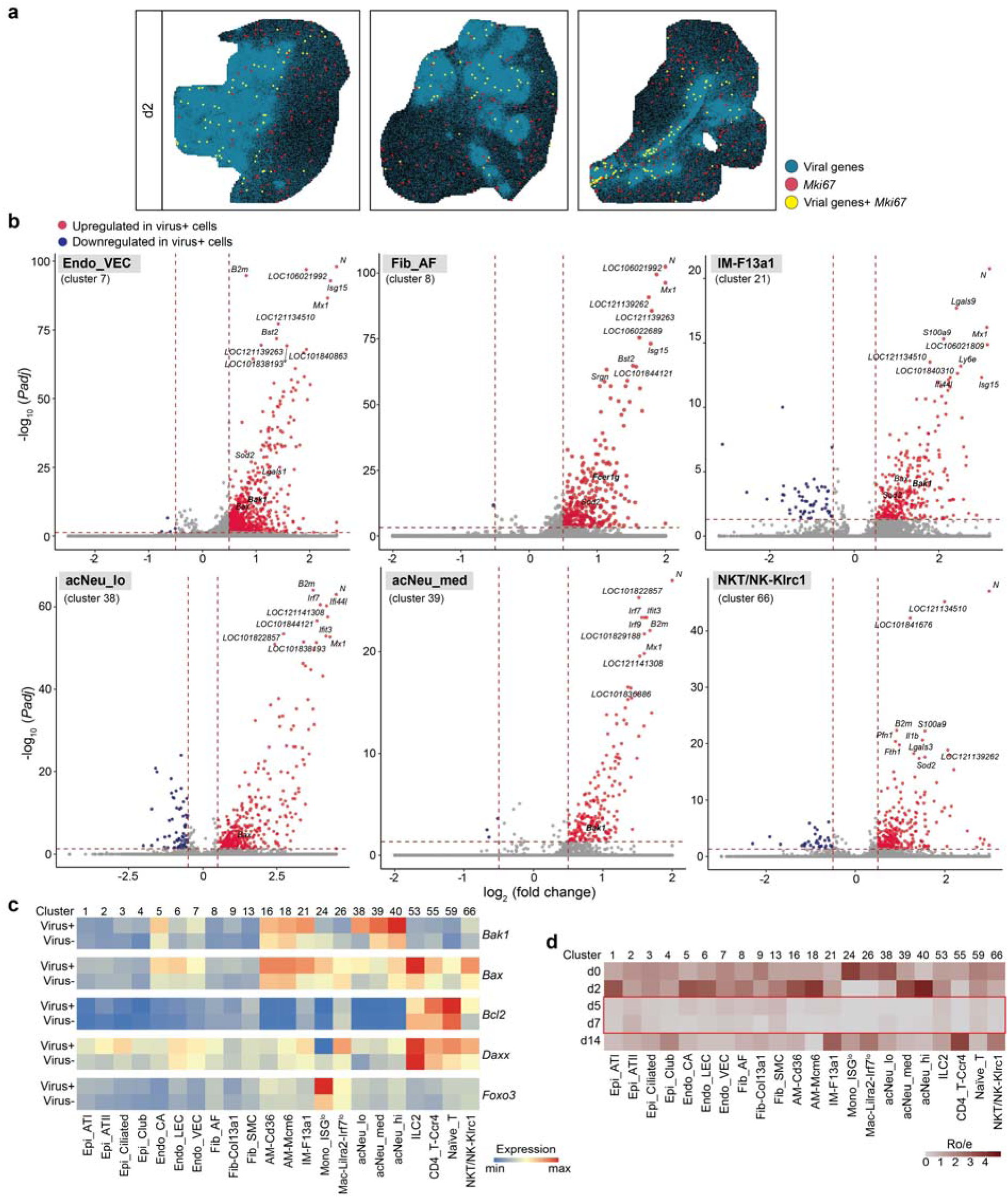
Acute damages to multiple cell types caused by SARS-CoV-2 infection. **a.** Spatial distribution of virus and AM-Cd36-Mki67 cells, represented by three lung sections at d2 (HD group). Magnified size of sparse spots was applied for clarity. **b.** Volcano plots of differentially expressed genes between virus-positive and virus-negative cells in representative cell subpopulations (cutoff: (|Log_2_FC|>0.5, *Padj*<0.05). **c.** Expression of cell death-related genes *Bak1*, *Bax*, *Bcl2*, *Daxx* and *Foxo3* in virus-positive and virus-negative cells of representative cell subpopulations. **d.** Temporal distribution of cell subpopulations in (**c**) that tended to be depleted after infection. Ro/e >1, enrichment; Ro/e<1, depletion.

**Extended Data Fig.5.**
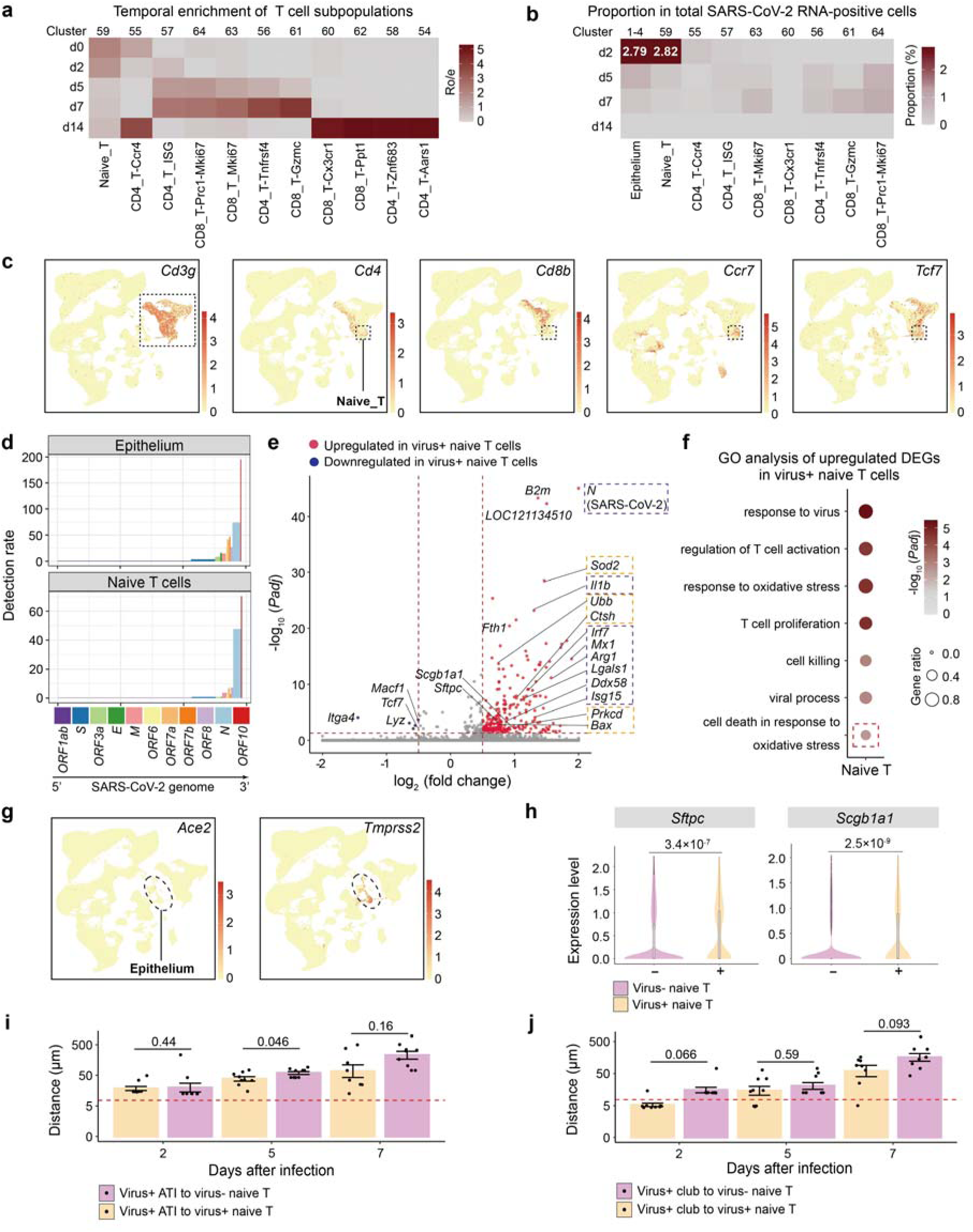
Signatures of lung naïve T cells invaded by SARS-CoV-2. **a.** Temporal distribution of T cell subpopulations. Ro/e >1, enrichment; Ro/e<1, depletion. **b.** Distribution of SARS-CoV-2 RNA-positive cells across epithelial cells and different T cell subpopulations at d2, d5, d7 and d14 (HD group). **c.** The expression of marker genes of T cells (*Cd3g, Cd4, Cd8b*) and marker genes of naïve T cells (*Ccr7*, *Tcf7*) in total lung cells. **d.** Detection rates of SARS-CoV-2 genes in epithelial cells and naïve T cells. **e.** Volcano plot of differentially expressed genes between virus-positive and virus-negative naïve T cells (cutoff: (|Log_2_FC|>0.5, *Padj*<0.05). Orange frame highlighted genes related to cell death. Gene description: *LOC121134510*, antileukoproteinase-like. **f.** GO terms enriched in upregulated genes in virus-positive naive T cells in **(e)**. **g.** The expression of *Ace2* and *Tmprss2* in total lung cells. **h.** The expression of ATII marker genes *Sftpc* and *Scgb1a1* in virus-positive naïve T cells quantified by log normalized expression. Two-sided Wilcoxon test. **i,j.** The peak values of the minimum distance between virus-positive ATI (**i**) or club cells (**j**) and their nearest virus-positive/negative naïve T cells of nine lung sections of each timepoint (HD group). Two-sided Student t-test.

**Extended Data Fig.6.**
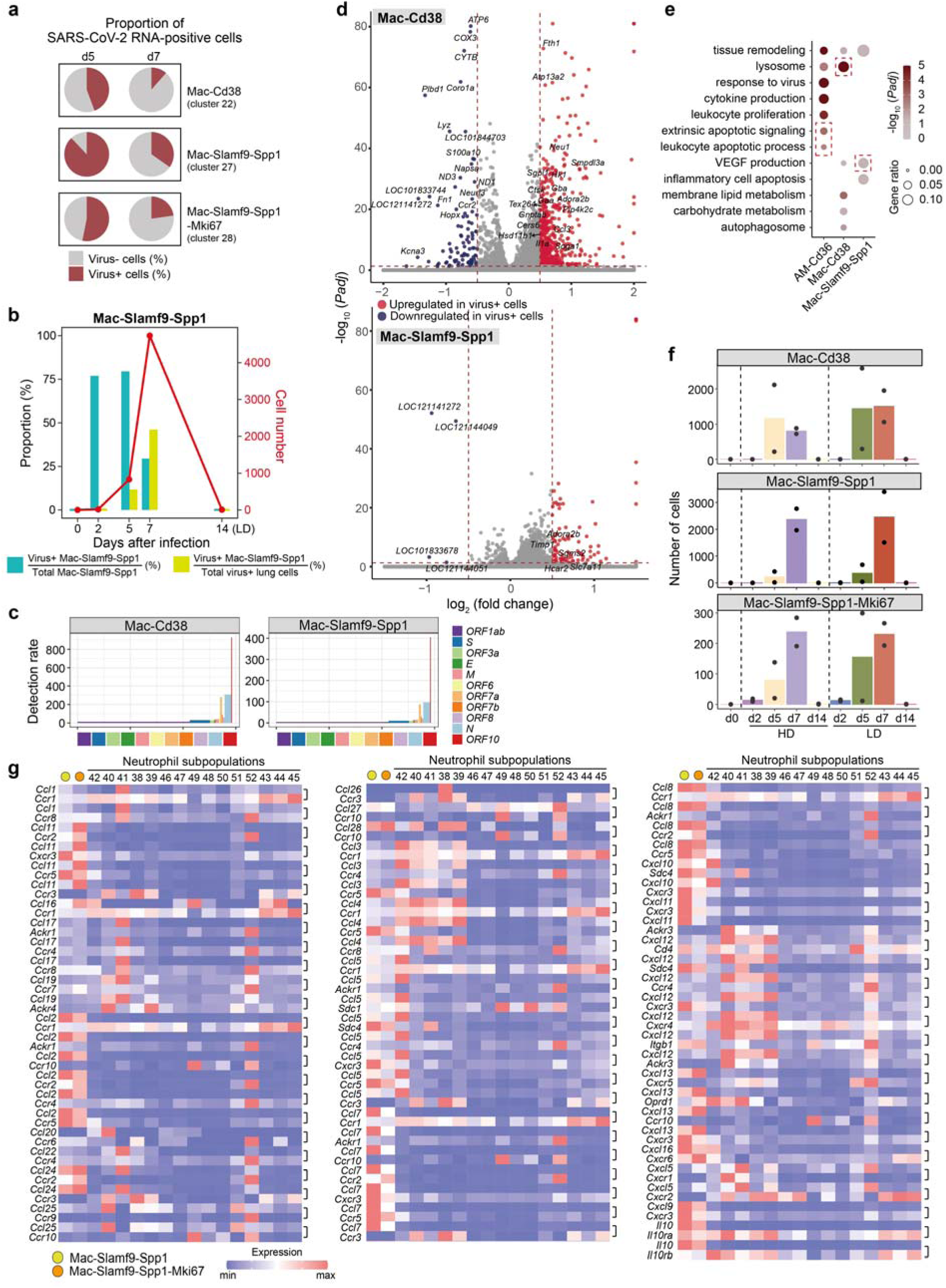
Signatures of *Cd38*^+^ and *Slamf9*^+^*Spp1*^+^ macrophages infected by SARS-CoV-2. **a.** The relative proportion of virus-positive cells within Mac-Cd38, Mac-Slamf9-Spp1 and Mac-Slamf9-Spp1-Mki67 subpopulations. More than 80% of Mac-Slamf9-Spp1 cells were virus-positive at d5. **b.** The number of total *Slamf9*^+^*Spp1*^+^ macrophages, the proportion of virus-positive *Slamf9*^+^*Spp1*^+^ macrophages and their proportion in total virus-positive cells at each timepoint (LD group). **c.** Detection rates of SARS-CoV-2 genes in *Cd38*^+^ and *Slamf9*^+^*Spp1*^+^ macrophages. **d.** Volcano plot of differentially expressed genes between virus-positive and virus-negative cells of *Cd38*^+^ and *Slamf9*^+^*Spp1*^+^ macrophages (cutoff: (|Log_2_FC|>0.5, *Padj*<0.05). **e.** GO terms enriched in upregulated genes in virus-positive AM-Cd36 (cluster 16), Mac-Cd38 cells (cluster 22) and Mac-Slamf9-Spp1 (cluster 27). **f.** Number of total cells of *Cd38*^+^, *Slamf9*^+^*Spp1*^+^ and proliferating *Slamf9*^+^*Spp1*^+^ macrophages detected at each timepoint. **g.** Expression of paired ligand and receptor genes in *Slamf9*^+^*Spp1*^+^, proliferating *Slamf9*^+^*Spp1*^+^ macrophages and neutrophil subpopulations.

**Extended Data Fig.7.**
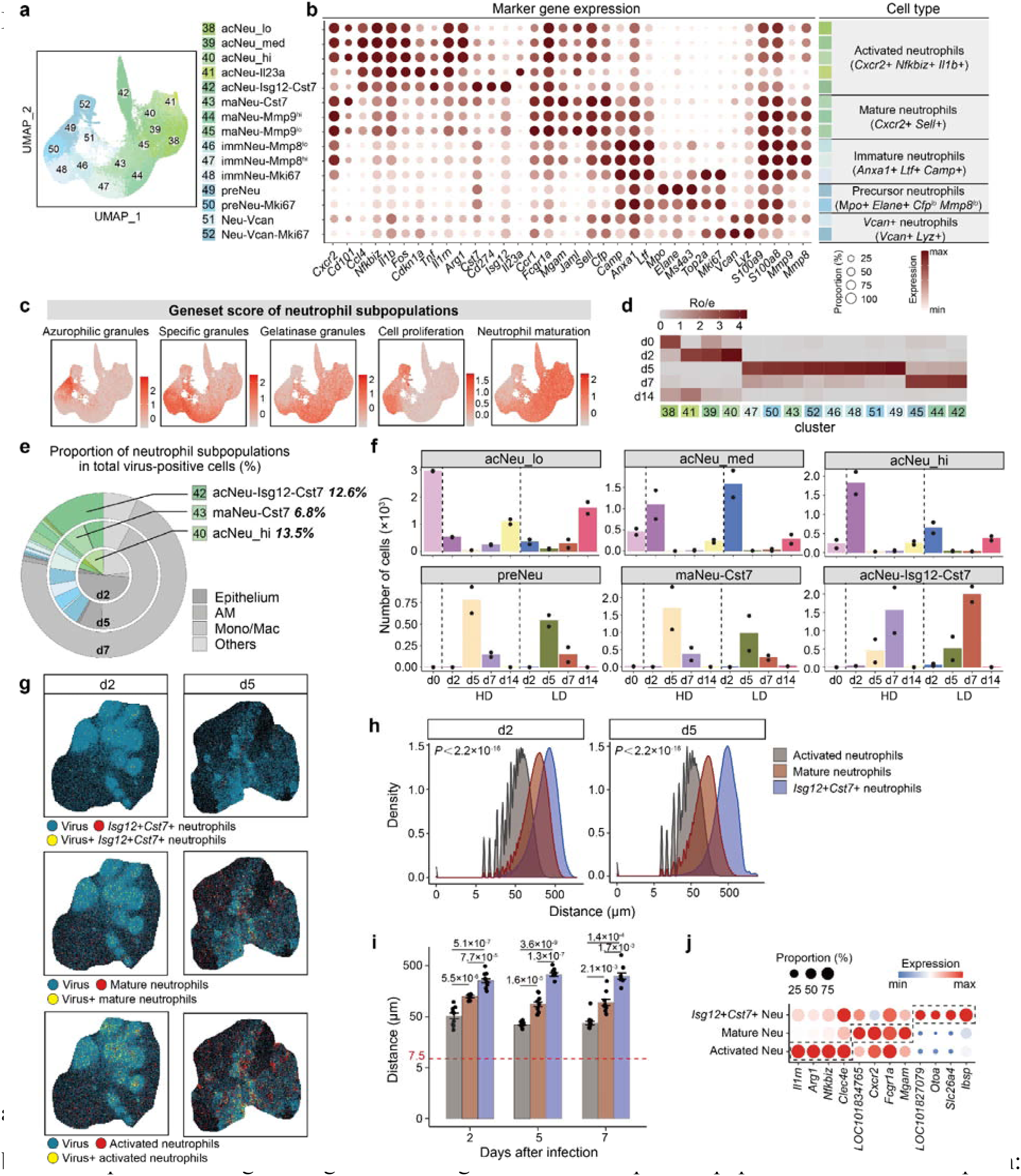
Neutrophil dynamics during SARS-CoV-2 infection. **a.** UMAP projection of neutrophils showing 15 cell subpopulations. **b.** The expression of signature genes defining different neutrophil subpopulations. Gene description : *LOC101831947, Ccl4; LOC101838467, Isg12; LOC101834765, Ccr1; LOC101825533, Cfp; LOC101829350, Camp; LOC101828213, S100a8*. **c.** Geneset score of cells in neutrophil subpopulations. **d.** Temporal distribution of neutrophil subpopulations. Ro/e >1, enrichment; Ro/e<1, depletion. **e.** The proportion of each cell subpopulation of virus-positive neutrophils in total virus-positive cells in d2, d5 and d7 (HD group). **f.** Number of total cells in key neutrophil subpopulations detected at each timepoint. **g.** Spatial distribution of virus and activated, mature or *Isg12^+^Cst7^+^* neutrophils, represented by a lung section at d2 and d5 (HD group). Magnified size of sparse spots was applied for clarity. **h.** Distance of viral spots to their nearest neutrophil subpopulations of nine lung sections of d2 and d5 (HD group) respectively. Kolmogorov-Smirnov test. **i.** The peak values of the minimum distance calculated in **(h)** of nine lung sections of each timepoint. Data are mean ± SEM. Two-sided Student t-test. **j.** The expression of genes that used to calculate spatial distances in the corresponding scRNA-seq neutrophil subpopulations.

**Extended Data Fig.8.**
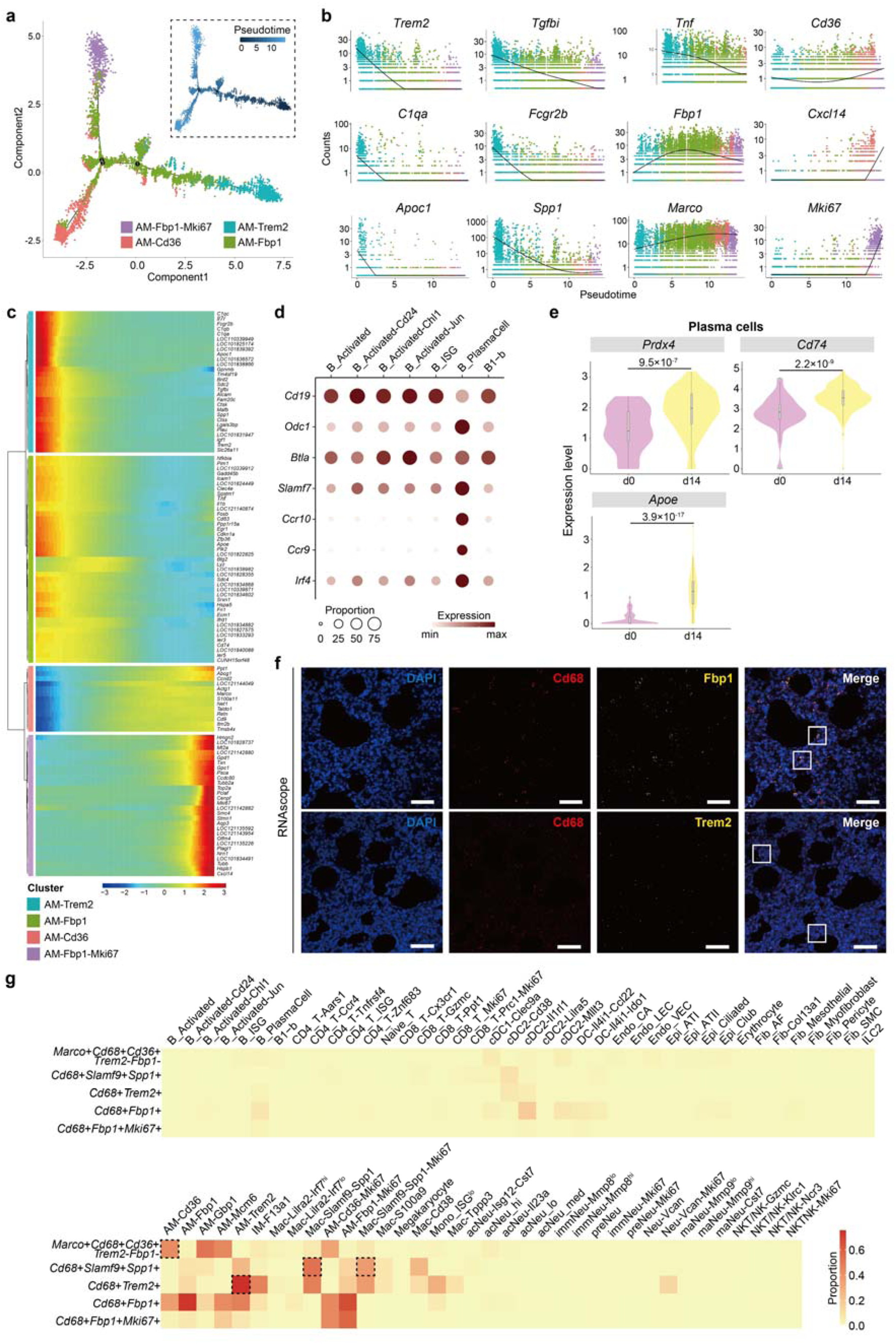
Signatures of *Fbp1*^+^ AMs, *Trem2*^+^ AMs and plasma cells for resolving inflammation. **a.** Pseudo-time trajectory initiated from AM-Trem2 towards AM-Fbp1, AM-Fbp1-Mki67 and AM-Cd36 predicted by Monocle2. **b.** Expression of representative genes in the route along the pseudo-time. **c.** Relative expression patterns of top 100 genes in the route along the pseudo time. **d.** The expression of signature genes of B/plasma cells. **e.** The expression of *Prdx4*, *Cd74* and *Apoe* in plasma cells of d0 and d14 quantified by log normalized expression. Two-sided Wilcoxon test. **f.** Staining of *Fpb1*^+^ AMs and *Trem2*^+^ AMs at d14 by RNAscope. Scale bar, 50μm. **g.** The expression of signature gens of *Slamf9*^+^*Spp1*^+^, *Trem2*^+^, *Fbp1*^+^, *Fbp1*^+^*Mki67*^+^, and *Marco*^+^*Cd36*^+^ macrophages in all 76 subpopulations identified in our study.

**Extended Data Fig.9.**
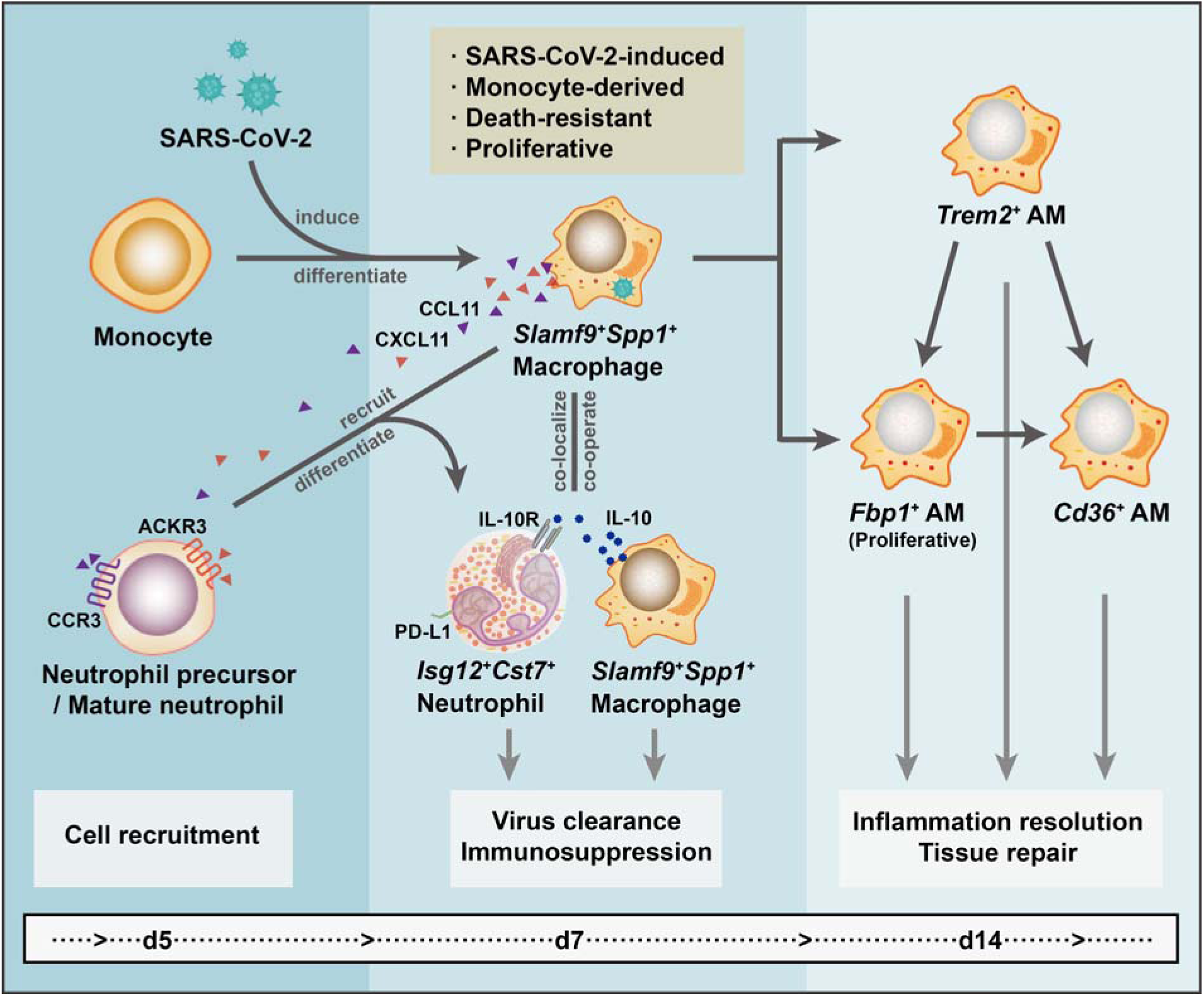
The dynamics and mechanisms of *Slamf9*^+^*Spp1*^+^ macrophages in clearing SARS-CoV-2 and resolving inflammation. *Slamf9*^+^*Spp1*^+^ macrophages are derived from peripheral monocytes, induced specifically by SARS-CoV-2 engulfment or infection in the lung, and are actively proliferating to win the arms race against SARS-CoV-2 intrahost transmission. These macrophages could secrete chemokines, such as *Ccl11* and *Cxcl11*, to recruit neutrophils and clear SARS-CoV-2 co-operatively. They could also produce IL-10 to induce an immunosuppressive niche and could be co-localized by *Isg12*^+^*Cst7*^+^ neutrophils to prevent inflammation-induced damages. After virus clearance, *Slamf9*^+^*Spp1*^+^ macrophages could transit into *Trem2*^+^, *Fbp1*^+^ and *Cd36*^+^ alveolar macrophages to further resolve inflammation and help for tissue repair.

## Methods

### Ethics Statement

At the Institute of Laboratory Animal Science (ILAS) of Chinese Academy of Medical Sciences, the animal biosafety level 3 (ABSL-3) facility was used to accomplish all the experiments with Syrian hamsters (male and female, aged 8-10 weeks) and hACE2 mice (male and female, aged 6-11 months). All experiments were implemented according to the Animal Welfare Act and other regulations associated with animals and experiments. The Institutional Animal Care and Use Committee (IACUC) of the ILAS, Peking Union Medical College & Chinese Academy of Medical Sciences, evaluated and gave permission to all the protocols in these studies, including animal experiments (Approval number QC21003).

### Viruses

The SARS-CoV-2 virus (accession number is MT093631.2, SARS CoV-2/WH-09/human/2020/CHN) was put into use in this study and propagated in Vero E6 cells incubated in Dulbecco’s modified Eagle’s medium (Invitrogen, Carlsbad, United States) supplemented with 10% fetal bovine serum (Gibco, Grand Island, United States) and incubated at 37 ◦C and 5% carbon dioxide.

### Experimental hamsters

Hamsters were intranasally inoculated with SARS-CoV-2 stock virus at 10^5^ 50% tissue culture infectious dose (TCID50) per mL (0.1 mL per animal) used as the high dose (HD) group, and 10^4^ TCID50 as the low dose (LD) group^1^. Hamsters intranasally inoculated with an equal volume of PBS were used as the healthy control (Ctrl) group. Hamsters were continuously monitored to record clinical symptoms, body weight, responses to external stimuli and death. Lung tissues of the infected group were collected at 2, 5, 7 and 14 days after infection (n=5/ timepoint), and those of the Ctrl group were collected at 2 days (n=5).

### Experimental hACE2 mice

Specific-pathogen-free, 6-11-month-old male and female hACE2 mice were obtained from the Institute of Laboratory Animal Sciences, Chinese Academy of Medical Sciences as described in our previous study^15^. The hACE2 mice were intranasally inoculated with SARS-CoV-2 stock virus at a dosage of 10^5^ TCID50, and those intranasally inoculated with an equal volume of PBS were used as healthy control (d0). The infected hACE2 mice were continuously observed to record body weight, clinical symptoms, responses to external stimuli and death. Mice were dissected at 7 and 14 days after infection (d7 and d14) to collect lung tissues for further experiments.

### Hematoxylin and Eosin (H&E) Staining

Hamster lungs were fixed in 10% formalin buffer saline (HT501128, Sigma-Aldrich) for 24 h at room temperature before paraffin embedding, and paraffin sections (4 µm in thickness) were applied following the routine operating procedure. The adjacent frozen sections (8-10 µm in thickness) of those sections used for Stereo-seq were fixed in acetone. All the tissue sections were stained with H&E. The pathological findings were carefully observed using an Olympus microscope. The pathological scorings of all hamster lungs were listed in **Supplementary Table 1**.

### *In situ* RNA hybridization

*In situ* RNA hybridization was performed using the Advanced Cell Diagnostics RNAscope Multiplex Fluorescent Detection kit v2 (323100, Bio-Techne) according to the manufacturer’s instructions. Staining of hamster lung specimens was performed using 3 μm paraffin-embedded thick sections. For multiplex staining the following probes were used: Fbp1 (Mau-Fbp1 1153521-C3), Cd4 (Mau-Cd4 1153531-C1), Il10 (Mau-Il10 1153551-C3), Mki67 (Mau-Mki67 1153561-C2), Trem2 (Mau-Trem2 1153571-C2), Sftpc (Mau-Sftpc 1058161-C3), Cd68 (Mau-Cd68 899591-C1), S (V-nCoV2019-S 848561-C2). Slides were counterstained with Mounting Medium With DAPI-Aqueous (ab104139, Abcam). Mounted slides were imaged on a Leica DMi8 fluorescent microscope (Leica Biosystems).

### Multiplex immunofluorescence staining

All collected lung tissues from hACE2 mice were fixed in 10% formalin buffer solution (HT501128, Sigma-Aldrich), and paraffin sections (3–4 µm in thickness) were prepared according to routine operating procedure^16,17^. The immunofluorescence staining was performed using a PANO IHC kit (cat. no. 0004100100; Panovue, Beijing, China) following the manufacturer’s instructions^18^. Different primary antibodies were sequentially applied to examine specific cell markers, including anti-CD68 Polyclonal antibody (28058-1-AP, 1:800; proteintech) and TREM2 Polyclonal antibody (13483-1-AP, 1:200; proteintech), or FBP1 Polyclonal antibody (12842-1-AP, 1:100, proteintech), or Osteopontin Polyclonal antibody (22952-1-AP, 1:200; proteintech) and SLAMF9 polyclonal antibody (LM-2205R, 1:200; LMAI BIO), followed by HRP-conjugated secondary antibody incubation and tyramide signal amplification (TSA). The slides were microwave-treated after each cycle of TSA. Nuclei were stained with 4’-6’-diamidino-2-phenylindole (DAPI; Sigma-Aldrich, St. Louis, MO) after antigen labeling. Stained slides were scanned using the 3D-histech (PANNORAMIC, 3DHISTECH, Hungary), which captures fluorescent spectra with identical exposure times using DAPI, FITC, TRICT and Cy5 channels, and the scans were combined to build a single stacked image.

### Single-cell RNA-seq library preparation

#### Single-cell suspension preparation

Fresh lung tissues were first rinsed with PBS, minced into small pieces by mechanical dissociation, and incubated for 1 h in 10 mL of DS-LT buffer (0.2 mg/mL CaCl_2_, 5 μM MgCl_2_, 0.2% BSA and 0.2 mg/mL Liberase in HBSS) at 37°C. After this, the tissue digestion was blocked by adding 3 mL of FBS, followed by filtration through a 100 µm cell strainer and centrifugation for 5 minutes at 500 *g* at 4°C. Samples were filtered through a 40 µm cell strainer and centrifuged for 5 minutes at 500 *g* at 4°C. Pellets were resuspended in cell resuspension buffer at 1,000 cells/μL for library preparation.

#### Library construction and sequencing

DNBelab C Series Single-Cell Library Prep Set (MGI, #1000021082) was used as described previously^2^. In brief, high-quality single-cell suspension was used for droplet generation, cell lysis and mRNA capture by microbeads were performed in the droplets, then emulsion breakage, beads collection, reverse transcription, finally the cDNA and droplet index were amplification to generate cDNA library and droplet index library respectively. All the processes were performed in the ABSL-3 laboratory till the PCR products of cDNA were gained. Library concentration was measured by use of Qubit™ ssDNA Assay Kit (Thermo Fisher Scientific, #Q10212) and sequenced by DNBSEQ T10 sequencer in the China National GeneBank (Shenzhen, China).

### Single-cell RNA-sequencing data processing

#### Raw data processing

For DNBelab C4 data, PISA (v0.7, https://github.com/shiquan/PISA) was used for sequencing reads filtering and demultiplexing. STAR (v2.7.9a)^3^ was applied to mapping reads with reference genome integrated by *Mesocricetus auratus* genome (BCM_Maur_2.0) and SARS-CoV-2 genome (NC_045512.2). UMI counting, cell identification, and generation of expression matrices were accomplished by PISA.

#### Single-cell RNA-seq data

The number of detected genes, the total UMI counts and proportion of mitochondrial gene counts per cell were used for quality control. Low quality cells with <1,000 UMI counts, <300 detected genes, or >5% mitochondrial gene counts were filtered. Cells with >60000 UMI counts or >10,000 detected genes were filtered out to remove potential doublets and multiplets. Then scDblFinder (v1.9.3) was applied to further identify potential doublets with its random methods (https://github.com/plger/scDblFinder).

#### Unsupervised cell clustering and annotation

Clustering analysis of the scRNA-seq dataset was performed using the Seurat package (v4.0.5) and the R program (v4.1.0). UMI count matrices of different samples were merged together for analysis, and then normalized (LogNormalization function, scale.factor=10000) and scaled with Seurat. 2,000 highly variable features were identified and a PCA matrix with 50 components was constructed. For clustering cells precisely, we applied two round clustering to the dataset. The first round clustering (resolution=0.4) identified approximation 5 major cell types including non-immune cells, myeloid cells, T cells, B cells and neutrophils. Clustering results were displayed by uniform manifold approximation and projection (UMAP)^4^ dimension reduction analysis. A second round of clustering was applied to each major cell type based on a procedure similar to the first round of clustering. Annotation of the final clusters was based on the known cell markers. Finally, 269,199 identified cells were remaining for downstream analysis. 76 clusters were finally obtained, representing different cell subpopulation. Additionally, we found that 31,924 cells in all cells were detected to have viral reads (UMI > 0). We compared cell numbers with viral genes at different days after infection. We also calculated detection rates of the 11 SARS-CoV-2 genes (*E*, *M*, *N*, *ORF10*, *ORF1ab*, *ORF3a*, *ORF6*, *ORF7a*, *ORF7b*, *ORF8* and *S*) along the viral genome from 5’ to 3’ according to a procedure previously described^5^ to exclude its relationship with library preparation methods. Additionally, Spearman’s correlation between the rate of virus-positive cells and the percentage of cluster acNeu-Isg12-Cst7 was calculated by the R function cor.test.

#### Gene module analysis

To evaluate neutrophil maturation and functional scores, we calculated the expression level of individual neutrophils with regarding predefined genesets (**Supplementary Table 8**)^6^. AddModuleScore function with default parameters was applied in this process. The resulting scores were shown in **Extended Data Fig.7c**.

#### Temporal distribution of different cell subpopulations

The temporal distribution of cell populations was indicated by Ro/e value, which was calculated by the following formula previously described (https://github.com/Japrin/STARTRAC)^7^:

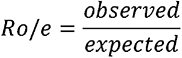

In which the observed variable is the cell number of a given *subpopulation_i_* at a specific *timepoint_j_*, and the expected variable is calculated by the following formula, which represents the expected cell number distribution for *subpopulation_i_* at *timepoint_j_*:

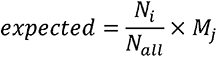

In which *N_all_* is the total cell number of all subpopulations, *N_i_* is the cell number of the given *subpopulation_i_* and *M_j_* is the cell number of all subpopulations at the specific *timepoint_j_*. *Role* > 1, it suggests that cells of the specific subpopulation are more frequently observed than random expectations at the specific timepoint (enrichment). If *Ro/e* < 1, it suggests that cells of the specific subpopulation are observed with less frequency than random expectations at the specific timepoint (depletion). The relevant results were shown in **Extended Data Fig.2d** and **Supplementary Table 5**.

#### Differential gene expression and Gene Ontology enrichment analysis

To investigate the effects of viral RNA in one subpopulation, we identified differentially expressed genes between viral RNA positive and negative cells by performing two-sided unpaired Wilcoxon tests based on the tl.rank_genes_groups function of the python package Scanpy (v1.8.1) (method=’wilcoxon’, corr_method = ‘benjamini-hochberg’). Differentially expressed genes were identified by two rules, *i.e.*, log_2_foldchanges >0.5 and P*adj*<0.05. Over-representation analysis were processed in 6 subpopulations (Naïve_T, AM-Cd36, Mac-Il10-Spp1, Mac-Cd38, acNeu-Isg12-Cst7 and acNeu_hi) using enrichGO (OrgDb=org.Mm.eg.db, pAdjustMethod=“BH”) function of the Cluster Profiler R package (v4.0.5)^8^. The relevant results were shown in **Supplementary Table 6**.

#### RNA velocity analysis

In order to find the relationships between different cell types in main lineages, RNA velocity analysis for neutrophils were conducted by using the scVelo python package^9^. The “basics” analysis was applied by following the tutorial (https://scvelo.readthedocs.io/VelocityBasics/). Different samples were analyzed separately and visualized based on the UMAP coordinates generated during the clustering step. In addition to neutrophils, monocytes/macrophages were analyzed independently according to the same procedure.

#### Pseudo-time analysis of macrophages

To analyze dynamic biological processes among AM-Trem2, AM-Fbp1, AM-Fbp1-Mki67 and AM-Cd36, we used the monocle R package (v2.20.0)^10–12^. We extracted UMI counts matrix of 4 subpopulation motioned above as input. We processed the pseudo-time analysis followed tutorial described in http://cole-trapnell-lab.github.io/monocle-release/docs/ with default parameters. Additionally, we sorted 517 genes out with parameters of mean_expression >= 0.1 and dispersion_empirical >= 1 * dispersion_fit for constructing single cell trajectories. After ordering cells, we clustered genes by pseudotemporal expression pattern. We got 110 genes belonging to four different clusters with statistical significance, which were showed in **Extended Data Fig.8c**.

#### Phenotypic linkage evaluation of myeloid cells between d7 and d14

We conducted cell type prediction analysis to scRNA-seq data of d7 and d14 (HD) cells to find potential phenotypic linkages between those cells. This process was performed by the functions FindTransferAnchors and TransferData function implemented in the software package Seurat, with 50 dims and all detected genes in scRNA-seq data as parameters. The prediction results of macrophages and monocytes were shown in **Fig.6a**.

#### Cell subpopulation similarity analysis between Syrian hamsters and COVID-19 patients

We also analyzed snRNA-seq data derived from autopsy lung tissues of COVID-19 patients^13^, to confirm whether subpopulations changes were similar between humans and Syrian hamsters. With homologene R package (v 1.4.68.19.3.27), we selected whole 13419 homologous genes, including *MARCO*/*Marco*, *CD68*/*Cd68*, *CD36*/*Cd36*, *TREM2*/*Trem2*, *FBP1*/*Fbp1*, *SLAMF9*/*Slamf9* and *MKI67*/*Mki67*. We applied a set of signature gene combinations to represent the cell types (Mac-Slamf9-Spp1: *CD68*^+^*SLAMF9*^+^*SPP1*^+^; AM-Trem2: *CD68*^+^*TREM2*^+^; AM-Fbp1: *CD68*^+^*FBP1*^+^; AM-Fbp1-Mki67: *CD68*^+^*FBP1*^+^*MKI67*^+^; AM-Cd36: *CD68*^+^*MARCO*^+^*CD36*^+^*TREM2*^-^*FBP1*^-^), which performed well in the scRNA-seq data generated by our study. Then we detected cell proportions with signature gene combinations expression, which were shown in **Fig.6h**.

### Stereo-seq library preparation and sequencing

#### Tissue processing

Lung tissue frozen sections were adhered to the Stereo-seq chip surface and incubated at 37L for 3-5 minutes. Then, the sections were fixed in methanol and incubated for 40 minutes at -20L before Stereo-seq library preparation. The same sections were stained with nucleic acid dye (Thermo fisher, Q10212) and imaging was performed with a Ti-7 Nikon Eclipse microscope prior to *in situ* capture at the channel of FITC.

#### Library construction and sequencing

Stereo-seq library construction was described previously^14^. For the Stereo-seq library construction of the Syrian hamster infected by SARS-CoV-2, all the processes were performed in the ABSL-3 laboratory till the PCR products of cDNA were gained. In brief, RNA released from the tissue was captured by the DNB on the chip after tissue sections were permeabilized, followed by reverse transcription. Then the cDNA was released from the stereo-chip and moved out of ABSL-3 lab for amplification. The concentration of PCR product was quantified by Qubit™ dsDNA Assay Kit (Thermo, Q32854). A total of 40∼60 ng of DNA was proceeded to fragmentation and amplification. PCR products were purified using the VAHTS^TM^ DNA Clean Beads (0.6× and 0.2×), used for DNB generation, and finally sequenced on MGI DNBSEQ-T10 sequencer.

#### Stereo-seq raw data processing

Fastq files were generated using a MGI DNBSEQ-T10 sequencer. *Mesocricetus auratus* genome (BCM_Maur_2.0) and SARS-CoV-2 genome were integrated as one reference for read mapping by STAR. One base mismatch was allowed to correct sequencing and PCR errors. Mapped reads were counted and annotated using an BGI-developed open-source pipeline SAW (https://github.com/BGIResearch/SAW).

#### Binning data of spatial Stereo-seq data

Because one Stereo-seq chip contained millions of DNBs (diameter: 220 nm), we merged adjacent 15*15 DNBs to one bin15 spot as a fundamental unit for downstream analysis, including cell type identification, spatial distance and cell density calculation. Additionally, matrices of different spatial Stereo-seq samples were normalized to 1,000,000 counts before analysis. For unsupervised clustering of the spatial transcriptomics data, bin80 spots were applied by merging adjacent 80*80 DNBs.

#### Quality control and batch effect correction of bin80 spatial Stereo-seq data

Stereo-seq data were quality-controlled by filtering low-quality spots for each chip based on the number of detected genes and the total UMI counts in each spot. We got 22,927 bin80 spots (40 μm resolution) per section, 557 genes and 1,115 UMI counts per spot of 15 histologically intact bin80 chips after quality control. Then bin80 Stereo-seq data matrices were merged, log-normalized and scaled by the R package Seurat, just like scRNA-seq matrices. We identified 2,000 highly variable features and calculated a PCA matrix with 20 components. RunHarmony function in the Harmony package was used to do batch effect correction for different chips.

#### Unsupervised spot clustering and subsets annotations

After unsupervised clustering (resolution=0.5) calculation, we identified 15 subsets of all selected bin80 spots of 15 selected chips. Clustering results were displayed by their spatial coordination (spatial_x and spatial_y) at the capture chip, which showed the real spatial orientation of spots in tissues. Annotation of the resulting clusters to tissue regions was based on the known markers and the spatial information. Subsets were named by the most enriched cell types or highly expressed genes of clusters (**Supplementary Table 2**).

#### Cell types identifying in bin15 spatial Stereo-seq data

Our bin15 spatial transcriptomics datasets contained an average of 656,311 bin15 spots (7.5 μm resolution) of all 81 chips. For 18 cell types we concerned, we used one to four marker genes of the corresponding cell types as surrogate markers (**Supplementary Table 3**) to investigate their spatial relationship with other cells. For SARS-CoV-2, bin15 spots with either of the 12 viral genes (*E*, *M*, *N*, *ORF10*, *ORF1a*, *ORF1ab*, *ORF3a*, *ORF6*, *ORF7a*, *ORF7b*, *ORF8* and *S*) positive were treated as virus-positive spots.

#### Distance between different cell types

With locations of spots containing cells we concerned, we calculated the minimum distance to other cells around. For each spot of cell type A, we calculated the Euclidean distance of this spot to its nearest spot of cell type B to represent their spatial relationship, based on the spatial coordinates of bin15 Stereo-seq data. Then we obtained a distance distribution for cell type A to cell type B. The peak value of the distance distribution in each chip was then extracted to summarize the spatial relationship from cell type A to cell type B. Obviously, this distance statistics was asymmetrical. Distance statistics from cell type A to cell type B and its reverse one were both used to interpret the spatial transcriptomics data.

#### Cell type density statistics

For each spot positive for marker genes of Mac-Slamf9-Spp1, AM-Fbp1 and death signals, we calculated the Gaussian density within its neighborhood for 18 other cell types. For spot i, whose coordinate is (*x^i^*,*y^i^*), the density was calculated by the following formula:

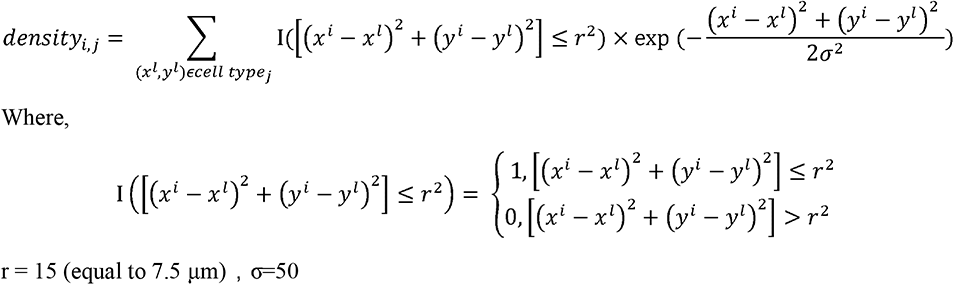

and the average Gaussian density in each chip of other 18 cell types in circle of different radii around death signals, Mac-Slamf9-Spp1 and M-Fbp1 were shown in **Fig.2j, 4c and 6d**.

### Statistical analysis

Statistics were processed with R statistical software v4.1.2 unless otherwise specified. Unpaired two-tailed t-tests, two-sided Wilcoxon test and Kolmogorov-Smirnov test were used as indicated.

## References

1. Sia, S. F. et al. Pathogenesis and transmission of SARS-CoV-2 in golden hamsters. Nature 583, 834–838, doi:10.1038/s41586-020-2342-5 (2020).

2. Du, S. et al. Structurally Resolved SARS-CoV-2 Antibody Shows High Efficacy in Severely Infected Hamsters and Provides a Potent Cocktail Pairing Strategy. Cell 183, 1013–1023 e1013, doi:10.1016/j.cell.2020.09.035 (2020).

3. Chen, A. et al. Spatiotemporal transcriptomic atlas of mouse organogenesis using DNA nanoball patterned arrays. Cell, in press (2022).

4. Qin, C. et al. Dysregulation of Immune Response in Patients With Coronavirus 2019 (COVID-19) in Wuhan, China. Clin Infect Dis 71, 762–768, doi:10.1093/cid/ciaa248 (2020).

5. Tan, M. et al. Immunopathological characteristics of coronavirus disease 2019 cases in Guangzhou, China. Immunology 160, 261–268, doi:10.1111/imm.13223 (2020).

6. Puelles, V. G. et al. Multiorgan and Renal Tropism of SARS-CoV-2. N Engl J Med 383, 590–592, doi:10.1056/NEJMc2011400 (2020).

7. Xu, Z. et al. Pathological findings of COVID-19 associated with acute respiratory distress syndrome. Lancet Respir Med 8, 420–422, doi:10.1016/S2213-2600(20)30076-X (2020).

8. Lowery, S. A., Sariol, A. & Perlman, S. Innate immune and inflammatory responses to SARS-CoV-2: Implications for COVID-19. Cell Host Microbe 29, 1052–1062, doi:10.1016/j.chom.2021.05.004 (2021).

9. Delorey, T. M. et al. COVID-19 tissue atlases reveal SARS-CoV-2 pathology and cellular targets. Nature 595, 107–113, doi:10.1038/s41586-021-03570-8 (2021).

10. Liao, M. et al. Single-cell landscape of bronchoalveolar immune cells in patients with COVID-19. Nat Med 26, 842–844, doi:10.1038/s41591-020-0901-9 (2020).

11. Zhang, J. Y. et al. Single-cell landscape of immunological responses in patients with COVID-19. Nat Immunol 21, 1107–1118, doi:10.1038/s41590-020-0762-x (2020).

12. Desai, N. et al. Temporal and spatial heterogeneity of host response to SARS-CoV-2 pulmonary infection. Nat Commun 11, 6319, doi:10.1038/s41467-020-20139-7 (2020).

13. Ren, X. et al. COVID-19 immune features revealed by a large-scale single-cell transcriptome atlas. Cell 184, 5838, doi:10.1016/j.cell.2021.10.023 (2021).

14. Rendeiro, A. F. et al. The spatial landscape of lung pathology during COVID-19 progression. Nature 593, 564–569, doi:10.1038/s41586-021-03475-6 (2021).

15. Cao, X. COVID-19: immunopathology and its implications for therapy. Nat Rev Immunol 20, 269–270, doi:10.1038/s41577-020-0308-3 (2020).

16. Song, Z. et al. SARS-CoV-2 Causes a Systemically Multiple Organs Damages and Dissemination in Hamsters. Front Microbiol 11, 618891, doi:10.3389/fmicb.2020.618891 (2020).

17. Rosenke, K. et al. Defining the Syrian hamster as a highly susceptible preclinical model for SARS-CoV-2 infection. Emerg Microbes Infect 9, 2673–2684, doi:10.1080/22221751.2020.1858177 (2020).

18. Yuan, S. et al. Clofazimine broadly inhibits coronaviruses including SARS-CoV-2. Nature 593, 418–423, doi:10.1038/s41586-021-03431-4 (2021).

19. Imai, M. et al. Syrian hamsters as a small animal model for SARS-CoV-2 infection and countermeasure development. Proc Natl Acad Sci U S A 117, 16587–16595, doi:10.1073/pnas.2009799117 (2020).

20. Hou, Y. J. et al. SARS-CoV-2 Reverse Genetics Reveals a Variable Infection Gradient in the Respiratory Tract. Cell 182, 429–446 e414, doi:10.1016/j.cell.2020.05.042 (2020).

21. Hoffmann, M. et al. SARS-CoV-2 Cell Entry Depends on ACE2 and TMPRSS2 and Is Blocked by a Clinically Proven Protease Inhibitor. Cell 181, 271–280 e278, doi:10.1016/j.cell.2020.02.052 (2020).

22. Jenkins, S. J. et al. Local macrophage proliferation, rather than recruitment from the blood, is a signature of TH2 inflammation. Science 332, 1284–1288, doi:10.1126/science.1204351 (2011).

23. Mathew, D. et al. Deep immune profiling of COVID-19 patients reveals distinct immunotypes with therapeutic implications. Science 369, doi:10.1126/science.abc8511 (2020).

24. Muus, C. et al. Single-cell meta-analysis of SARS-CoV-2 entry genes across tissues and demographics. Nat Med 27, 546–559, doi:10.1038/s41591-020-01227-z (2021).

25. Amici, S. A. et al. CD38 Is Robustly Induced in Human Macrophages and Monocytes in Inflammatory Conditions. Front Immunol 9, 1593, doi:10.3389/fimmu.2018.01593 (2018).

26. Zeidler, J. D., Kashyap, S., Hogan, K. A. & Chini, E. N. Implications of the NADase CD38 in COVID pathophysiology. Physiol Rev 102, 339–341, doi:10.1152/physrev.00007.2021 (2022).

27. Veillette, A. Immune regulation by SLAM family receptors and SAP-related adaptors. Nat Rev Immunol 6, 56–66, doi:10.1038/nri1761 (2006).

28. Dragovich, M. A. & Mor, A. The SLAM family receptors: Potential therapeutic targets for inflammatory and autoimmune diseases. Autoimmun Rev 17, 674–682, doi:10.1016/j.autrev.2018.01.018 (2018).

29. Morse, C. et al. Proliferating SPP1/MERTK-expressing macrophages in idiopathic pulmonary fibrosis. Eur Respir J 54, doi:10.1183/13993003.02441-2018 (2019).

30. La Manno, G. et al. RNA velocity of single cells. Nature 560, 494–498, doi:10.1038/s41586-018-0414-6 (2018).

31. Gauthier, T. & Chen, W. Modulation of Macrophage Immunometabolism: A New Approach to Fight Infections. Front Immunol 13, 780839, doi:10.3389/fimmu.2022.780839 (2022).

32. Liu, Y. et al. N (6)-methyladenosine RNA modification-mediated cellular metabolism rewiring inhibits viral replication. Science 365, 1171–1176, doi:10.1126/science.aax4468 (2019).

33. Filep, J. G. & Ariel, A. Neutrophil heterogeneity and fate in inflamed tissues: implications for the resolution of inflammation. Am J Physiol Cell Physiol 319, C510–C532, doi:10.1152/ajpcell.00181.2020 (2020).

34. Perisic Nanut, M., Sabotic, J., Svajger, U., Jewett, A. & Kos, J. Cystatin F Affects Natural Killer Cell Cytotoxicity. Front Immunol 8, 1459, doi:10.3389/fimmu.2017.01459 (2017).

35. Li, X. et al. High level expression of ISG12(1) promotes cell apoptosis via mitochondrial-dependent pathway and so as to hinder Newcastle disease virus replication. Vet Microbiol 228, 147–156, doi:10.1016/j.vetmic.2018.11.017 (2019).

36. Ullah, H. et al. Antiviral Activity of Interferon Alpha-Inducible Protein 27 Against Hepatitis B Virus Gene Expression and Replication. Front Microbiol 12, 656353, doi:10.3389/fmicb.2021.656353 (2021).

37. Rossaint, J. et al. Directed transport of neutrophil-derived extracellular vesicles enables platelet-mediated innate immune response. Nat Commun 7, 13464, doi:10.1038/ncomms13464 (2016).

38. Henn, D. et al. Xenogeneic skin transplantation promotes angiogenesis and tissue regeneration through activated Trem2(+) macrophages. Sci Adv 7, eabi4528, doi:10.1126/sciadv.abi4528 (2021).

39. Geiger, R. et al. L-Arginine Modulates T Cell Metabolism and Enhances Survival and Anti-tumor Activity. Cell 167, 829–842 e813, doi:10.1016/j.cell.2016.09.031 (2016).

40. Wang, S. et al. A single-cell transcriptomic landscape of the lungs of patients with COVID-19. Nat Cell Biol 23, 1314–1328, doi:10.1038/s41556-021-00796-6 (2021).

41. Sanjiv, N., Osathanugrah, P., Fraser, E., Ng, T. F. & Taylor, A. W. Extracellular Soluble Membranes from Retinal Pigment Epithelial Cells Mediate Apoptosis in Macrophages. Cells 10, doi:10.3390/cells10051193 (2021).

42. Schulte-Schrepping, J. et al. Severe COVID-19 Is Marked by a Dysregulated Myeloid Cell Compartment. Cell 182, 1419–1440 e1423, doi:10.1016/j.cell.2020.08.001 (2020).

43. Lopez-Sampalo, A., Bernal-Lopez, M. R. & Gomez-Huelgas, R. Persistent COVID-19 syndrome. A narrative review. Rev Clin Esp (Barc), doi:10.1016/j.rceng.2021.10.001 (2022).

## References

1. Song, Z. et al. SARS-CoV-2 Causes a Systemically Multiple Organs Damages and Dissemination in Hamsters. Front Microbiol 11, 618891, doi:10.3389/fmicb.2020.618891 (2020).

2. Zhu, L. et al. Single-Cell Sequencing of Peripheral Mononuclear Cells Reveals Distinct Immune Response Landscapes of COVID-19 and Influenza Patients. Immunity 53, 685–696 e683, doi:10.1016/j.immuni.2020.07.009 (2020).

3. Dobin, A. et al. STAR: ultrafast universal RNA-seq aligner. Bioinformatics 29, 15–21, doi:10.1093/bioinformatics/bts635 (2013).

4. Becht, E. et al. Dimensionality reduction for visualizing single-cell data using UMAP. Nat Biotechnol, doi:10.1038/nbt.4314 (2018).

5. Ren, X. et al. COVID-19 immune features revealed by a large-scale single-cell transcriptome atlas. Cell 184, 5838, doi:10.1016/j.cell.2021.10.023 (2021).

6. Xie, X. et al. Single-cell transcriptome profiling reveals neutrophil heterogeneity in homeostasis and infection. Nat Immunol 21, 1119–1133, doi:10.1038/s41590-020-0736-z (2020).

7. Zhang, L. et al. Lineage tracking reveals dynamic relationships of T cells in colorectal cancer. Nature 564, 268–272, doi:10.1038/s41586-018-0694-x (2018).

8. Yu, G., Wang, L. G., Han, Y. & He, Q. Y. clusterProfiler: an R package for comparing biological themes among gene clusters. OMICS 16, 284–287, doi:10.1089/omi.2011.0118 (2012).

9. Bergen, V., Lange, M., Peidli, S., Wolf, F. A. & Theis, F. J. Generalizing RNA velocity to transient cell states through dynamical modeling. Nat Biotechnol 38, 1408–1414, doi:10.1038/s41587-020-0591-3 (2020).

10. Trapnell, C. et al. The dynamics and regulators of cell fate decisions are revealed by pseudotemporal ordering of single cells. Nat Biotechnol 32, 381–386, doi:10.1038/nbt.2859 (2014).

11. Qiu, X. et al. Single-cell mRNA quantification and differential analysis with Census. Nat Methods 14, 309–315, doi:10.1038/nmeth.4150 (2017).

12. Qiu, X. et al. Reversed graph embedding resolves complex single-cell trajectories. Nat Methods 14, 979–982, doi:10.1038/nmeth.4402 (2017).

13. Wang, S. et al. A single-cell transcriptomic landscape of the lungs of patients with COVID-19. Nat Cell Biol 23, 1314–1328, doi:10.1038/s41556-021-00796-6 (2021).

14. Chen, A. et al. Spatiotemporal transcriptomic atlas of mouse organogenesis using DNA nanoball patterned arrays. Cell, accepted.

15. Bao, L. et al. The pathogenicity of SARS-CoV-2 in hACE2 transgenic mice. Nature 583, 830–833, doi:10.1038/s41586-020-2312-y (2020).

16. Yu, P. et al. Age-related rhesus macaque models of COVID-19. Animal Model Exp Med 3, 93–97, doi:10.1002/ame2.12108 (2020).

17. Ma, Y. et al. SARS-CoV-2 infection aggravates chronic comorbidities of cardiovascular diseases and diabetes in mice. Animal Model Exp Med 4, 2–15, doi:10.1002/ame2.12155 (2021).

18. Song, Z. et al. Integrated histopathological, lipidomic, and metabolomic profiles reveal mink is a useful animal model to mimic the pathogenicity of severe COVID-19 patients. Signal Transduct Target Ther 7, 29, doi:10.1038/s41392-022-00891-6 (2022).

